# *Histomonas meleagridis* infection and vaccination protection in chickens – a metabolomics study

**DOI:** 10.64898/2026.09.24.753691

**Authors:** Anh Vu Nguyen, Topi Meuronen, Atte Lihtamo, Soile Turunen, Tamas Hatfaludi, Michael Hess, Antton Alberdi, Kati Hanhineva, Olli Kärkkäinen

## Abstract

Histomonosis, caused by *Histomonas meleagridis*, is a severe disease in poultry with main pathological lesions in ceca and liver, leading to significant animal suffering and economic losses, with no licensed treatments available. While cecal pathology and microbiota disruption have been documented, the metabolic consequences of infection and vaccination remain poorly understood. Here, we used untargeted LC-MS metabolomics to profile cecal tissue and luminal digesta in a controlled vaccination-challenge chicken trial (n=100, 25 birds/group), including uninfected and unvaccinated but challenged controls, together with birds vaccinated either cloacally, or orally via an edible gel following challenge, across five post-challenge timepoints. Results showed that *H. meleagridis* infection caused extensive, time-dependent disruption of the cecal metabolome, especially at 7-14 days post-infection (DPI). At DPI 14, 325 tissue and 359 digesta metabolites were significantly altered in unvaccinated challenged birds. Infection was associated with depletion of microbiota-related metabolites, including dicarboxylic acids, tryptophan indoles, phytochemical derivatives, bilirubin catabolites, phenolics, vitamins, and secondary bile acids, alongside accumulation of microbial substrates and fermentation products. Changes in the host-associated metabolites indicated mucosal inflammation, epithelial injury, mitochondrial and amino acid metabolic disturbance, oxidative stress, and membrane lipid remodeling. Cloacal vaccination provided greater metabolic protection than oral vaccination, with 78.3% of disrupted metabolites preserved at DPI 14, compared with 60.7% vaccination by oral gel delivery, pointing towards the importance of vaccine take. By DPI 21, cloacally vaccinated birds showed nearly complete metabolic recovery across most pathway groups, whereas gel-vaccinated birds retained broader metabolic disturbance. These findings reveal histomonosis as a disruption of the cecal host-microbiota metabolic interface and provide a metabolite-level framework for future studies of pathogenesis, vaccine development, and prevention strategies.

## 1. Introduction

*Histomonas meleagridis* is an extracellular protozoan flagellate and causative agent of histomonosis, a necrotizing disease of the ceca and liver first described in domestic turkeys in 1893 (Cushman, 1893). In susceptible turkeys, outbreaks can cause mortalities approaching 100%, whereas in chickens infection causes lower mortality but results in cecal inflammation, impaired performance, and reduced egg production (Lesleigh C. Beer et al., 2022; Hess et al., 2015; McDougald, 2005). Furthermore, surviving birds may act as reservoirs for onward transmission. For much of the twentieth century, histomonosis was effectively controlled by chemoprophylactics, including arsenicals, nitroimidazoles, and nitrofurans, but these compounds were progressively withdrawn from European and North American markets between the mid-1990s and 2015s due to safety concerns (Clark and Kimminau, 2017; Liebhart et al., 2017). Since then, the disease has re-emerged as a major challenge for poultry production in the absence of licensed drugs or vaccines, creating an urgent need to understand the biological processes that determine disease severity and protection to inform new treatment strategies (Landim de Barros et al., 2022; Liebhart and Hess, 2020).

The cecum lies at the center of histomonosis pathogenesis. It is the main site of parasite-associated lesions and one of the most densely microbial colonized compartments of the avian gut, with complex communities that supply metabolites essential for mucosal homeostasis (Ali et al., 2022). Previous studies have shown that *H. meleagridis* infection markedly disrupts the cecal microbiota, reducing richness and diversity, shifting dominant bacterial phyla, and depleting commensal groups such as Ruminococcaceae and Lactobacillaceae (Abdelhamid et al., 2021, 2020; Rafieian-Naeini et al., 2025). Experimentally induced typhlitis characterized by severe mucosal lesions coinciding with clinical signs and production losses is most evident during the first two weeks post-infection (Hauck and Hafez, 2013). Experimental live-attenuated vaccination has emerged as a promising control strategy, and both cloacal and oral delivery can protect against challenge in controlled trials, although efficacy is strongly influenced by age, route, and formulation (L. C. Beer et al., 2022; Chen et al., 2024; Mitra et al., 2018). However, little is known about how infection and vaccination reshape the local biochemical environment of the cecum and which metabolic pathways distinguish destructive from protective responses.

To date, metabolomics investigations of histomonosis are limited. One recent study characterized the metabolome of *H. meleagridis in vitro*, highlighting dependencies on bacterial purine and riboflavin supply, while another ^1^H-NMR-based analysis examined systemic plasma and liver metabolite profiles in a concurrent nematode-histomonas infection model (Ammar et al., 2024; Oladosu et al., 2023). These studies, however, did not resolve the *in vivo* metabolic changes occurring directly in cecal tissue and luminal contents during *H. meleagridis* infection, nor how these changes are modulated by vaccination. Thus, the metabolic pathways linking cecal dysbiosis, mucosal damage, and vaccine-mediated protection remain largely undefined.

In this study, we aimed to comprehensively characterize the cecal metabolic responses to *H. meleagridis* infection in chickens and to understand the modulatory effects of different vaccination routes using untargeted LC-MS metabolomics. We profiled cecal tissue and digesta at five time points post-challenge in a controlled vaccination trial, including uninfected controls, and infected birds with cloacal, oral vaccination via an edible gel, or unvaccinated challenges as reported previously (Hatfaludi et al., in prep.). This enabled us to resolve microbiota- and host-associated metabolic signatures of histomonosis and their modulation by vaccination route. This study provides the first metabolite-level framework for understanding histomonosis pathogenesis in the cecum of chickens and identifies metabolic features associated with vaccination-mediated protection, offering new insights into the pathogenesis of histomonosis and related enteric infections with implications for vaccine design.

## 2. Materials and Methods

### 2.1. Study design and sample collection

The cecal metabolome of chickens was characterized from samples collected during a recently reported controlled *Histomonas meleagridis* vaccination/challenge trial (Hatfaludi et al., in prep.). Briefly, one hundred specific pathogen-free (SPF) chickens were assigned to four experimental groups (n = 25 per group): T1 (cloacal vaccination/challenge), T2 (oral vaccination/challenge), T3 (-/challenge), and T4 (-/-), corresponding to cloacal vaccination followed by *H. meleagridis* challenge; oral vaccination by edible gel followed by challenge; challenge without prior vaccination, infected control; and unvaccinated, unchallenged, negative control (**Fig. 1**). Vaccination was administered at 7 days of age using an *in vitro* attenuated clonal monoxenic culture of *H. meleagridis*/Turkey/Austria/2922-C6/04-4CEF (passage 351). T1 birds received a cloacal dose of **∼** 10^4^ cells, while T2 birds received a calculated oral dose of ∼ 2×10^5^ cells within a gel formulation. At 35 days of age, groups T1, T2, and T3 were challenged with the virulent strain *H. meleagridis*/Turkey/Germany/4114-C18/05 (passage 8). Five birds each per group were sacrificed at DPI 0 (pre-challenge baseline) and four timepoints post-challenge: DPI 4, DPI 7, DPI 14, and DPI 21, yielding 100 cecal tissue samples and 98 cecal digesta samples (one T3 digesta sample at DPI 21 and one T2 digesta sample at DPI 7 were unavailable). All birds survived throughout the 21-day post-challenge period, enabling complete longitudinal sampling across all groups.

**Fig. 1.**
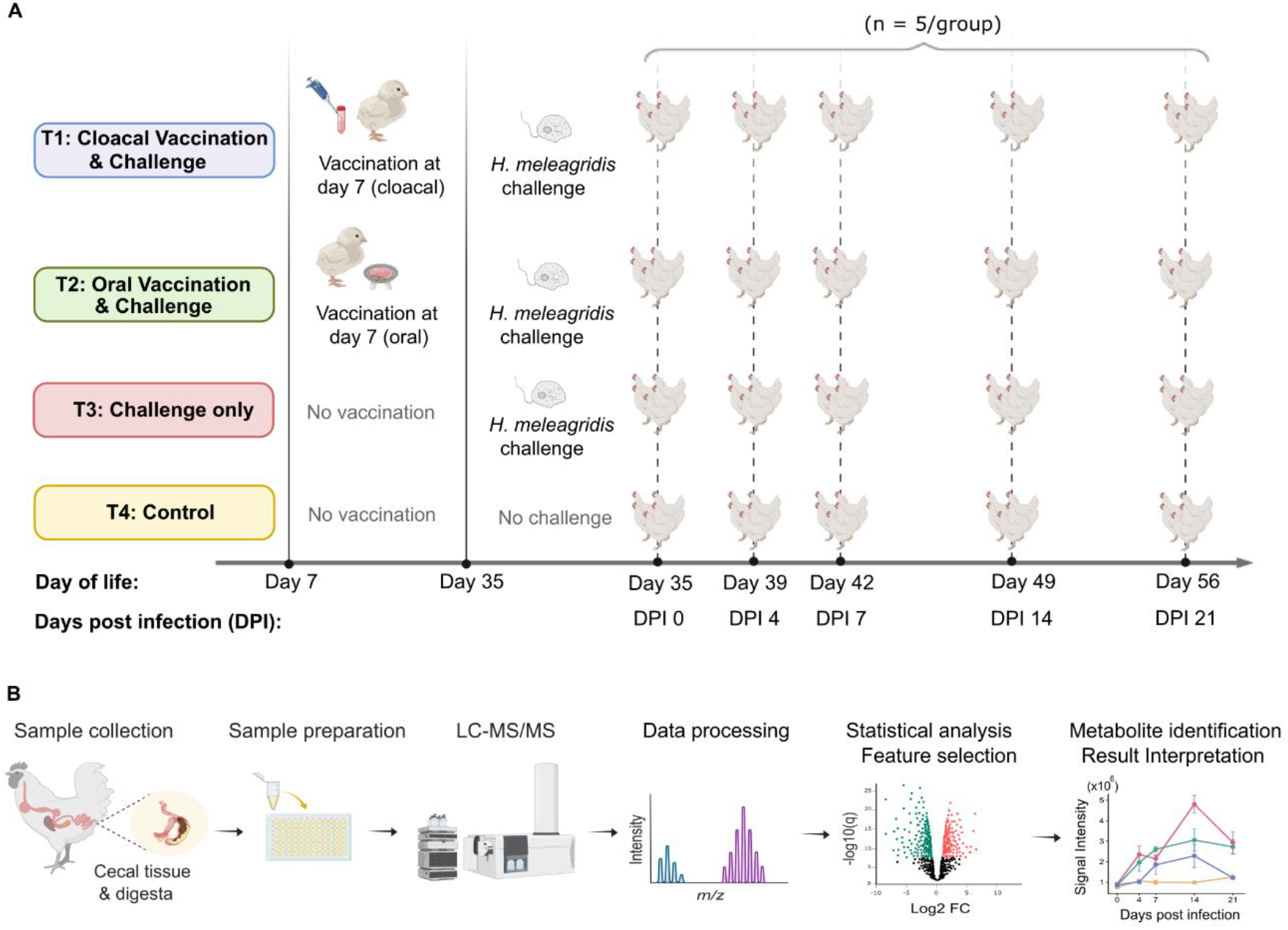
Study design and untargeted LC-MS metabolomics workflow. **(A)** One hundred SPF chickens were assigned to four groups (n = 25/group): T1 (cloacal vaccination/challenge), T2 (oral vaccination/challenge), T3 (-/challenge), and T4 (-/-). Vaccination was administered at day 7 of life and challenge with a virulent *H. meleagridis* strain at day 35 (DPI 0). Five birds per group were sacrificed at DPI 0, 4, 7, 14, and 21, and cecal tissue and digesta samples were collected for metabolomics analysis. **(B)** Cecal tissue and digesta samples underwent solvent-based extraction and filtration (Sample preparation), followed by four-mode LC-MS/MS analysis combining reversed-phase (RP) chromatography and hydrophilic interaction liquid chromatography (HILIC) in positive and negative ionization modes. Raw data were processed in MS-DIAL and preprocessed in R using *notame* workflow (Koistinen et al., 2026), including drift correction, quality filtering, and probabilistic quotient normalization (Data processing). Differentially abundant features were identified by statistical analysis and multivariate feature selection (Statistical analysis, Feature selection). Annotated features were manually curated before the identified metabolites were interpreted further (Metabolite identification, Result interpretation). The figure was created with Biorender.com.

### 2.2. Sample preparation

Frozen cecal tissue and digesta samples were homogenized by bead mill at 0**-**4 °C and extracted using 80% methanol (v/v) at standardized sample-to-solvent ratios (1000 µL per 100 mg tissue; 1500 µL per 100 mg digesta). Extracts were centrifuged at 17,000 × g, filtered through 0.2 µm filter plates, and stored at 2**-**8 **°**C until analysis. Pooled quality control (QC) samples were prepared by combining equal aliquots from all supernatants of each sample type and were used for instrument conditioning, drift correction, and quality monitoring throughout the analytical run. Full extraction details are provided in the **Supplementary Method**.

### 2.3. LC-MS analysis

Liquid chromatography-mass spectrometry (LC-MS) analysis was performed on an Agilent 1290 Infinity II UHPLC coupled to an Agilent 6546 QTOF mass spectrometer with a Dual Jet Stream electrospray ionization (ESI) source (Avella et al., 2026). To maximize metabolome coverage, each sample was analyzed in complementary analytical modes: reversed-phase (RP) chromatography and hydrophilic interaction liquid chromatography (HILIC) in both positive and negative ionization modes. MS/MS data were acquired by data-dependent acquisition (DDA) at collision energies of 10, 20, and 40 eV. Raw data were processed in MS-DIAL (*v4.90*) and subsequently imported into R for preprocessing using the *notame* package (Koistinen et al., 2026), including blank filtering, drift correction by cubic spline fitting to pooled QC injections, and feature quality filtering based on robust RSD and D-ratio thresholds. Between-sample normalization used probabilistic quotient normalization (PQN). All samples were measured within a single analytical batch. Full LC-MS acquisition and preprocessing parameters are provided in the **Supplementary Method**.

### 2.4. Metabolite identification

Putative identifications were assigned by automated matching against in-house and public spectral libraries (HMDB, LipidMaps, KEGG, ChEBI) based on accurate mass, retention time, and MS/MS spectral similarity, followed by manual curation. Identification confidence was assigned according to the Metabolomics Standards Initiative (MSI) framework (Sumner et al., 2007): level 1 (confirmed by authentic reference standard analyzed under the same conditions; MS1 ≤ 0.01 Da, MS2 ≤ 0.05 Da, RT ≤ 0.3 min), level 2 (putatively annotated by spectral library match with MS/MS confirmation; MS1 ≤ 0.01 Da, MS2 ≤ 0.05 Da), and level 3 (putatively characterized by accurate mass and molecular formula assignment using MS-FINDER v3.73; MS1 ≤ 0.01 Da). Where a metabolite was detected across multiple analytical modes, a single representative feature was selected based on signal quality; redundant cross-mode features were excluded.

### 2.5. Statistical analysis

Feature intensities were log_2_-transformed (log_2_(x+1)) prior to modeling. To resolve treatment effects at each sampling point independently and avoid imposing parametric assumptions on the temporal response, statistical models were fitted separately for each combination of time point and sample type, yielding ten independent models (5 DPI × 2 sample types). Within each model, differential abundance was assessed using empirical Bayes linear modeling (*limma*) with formula ∼TreatmentID and T4 as reference, providing three contrasts: T3 vs T4, T1 vs T4, and T2 vs T4. Multiple testing correction used the Benjamini-Hochberg procedure. Metabolites with FDR-adj.p (q) < 0.05 were considered statistically significant. Effect sizes are reported as log_2_ fold change (log_2_FC) or FC relative to T4. The proportion of metabolome variance explained by treatment group at each time point was quantified by PERMANOVA (*vegan::adonis2*, 999 permutations, marginal effects). To assess vaccination-mediated metabolic protection, each metabolite significantly disrupted by infection (T3 vs T4, adj.p < 0.05) was classified as protected if the corresponding vaccinated group (T1 or T2) was not significantly different from T4 (adj.p ≥ 0.05), indicating restoration towards healthy control levels, or as unprotected if it remained significantly altered in the same direction as T3.

### 2.6. Data visualization

All visualizations were produced in R (*v4.5.2*); volcano plots, score plots, and line plots used ggplot2, and heatmaps used ComplexHeatmap. Sample-level metabolome structure was explored by principal component analysis (PCA; *pcaMethods::pca*, unit-variance scaling). Model quality was assessed by R^2^ (goodness-of-fit) and Q^2^ (predictive power estimated by 7-fold cross-validation). Differential abundance results were visualized as volcano plots of log_2_FC versus - log_10_(adj.p) for each timepoint and contrast, with a shared y-axis within each sample type to enable direct temporal comparison. Metabolite abundance patterns were summaried using a heatmap of all identified metabolites (ComplexHeatmap; row-scaled Z-scores of group means across all treatment × timepoint conditions). Longitudinal abundance profiles (mean ± SEM per treatment across DPI) were plotted for representative metabolites.

## 3. Results

### 3.1. *H. meleagridis* infection broadly disrupts the cecal metabolome

We performed LC-MS profiling across four analytical modes (HILIC± and RP±). After merging preprocessed matrices from four analytical modes, 25901 and 27560 features passed quality filtering from cecal tissue and cecal digesta, respectively; 35844 high-quality features were detected in at least one intestinal compartment and 17617 in both simultaneously (**Table S1**). After manual curation, we identified 555 metabolites that significantly changed compared to the unchallenged control (T4) in at least one treatment, compartment or timepoint (**Table S2**). PCA of the full dataset revealed broad separation of samples by both treatment groups and days post-infection (DPI) in both cecal tissue and digesta samples (**Fig. 2A-B**). In both compartments, the cloacally vaccinated group (T1) tended to cluster closer to the control, whereas the orally vaccinated group (T2) more often clustered closer to the infected unvaccinated group (T3), suggesting a stronger overall preservation of a control-like metabolic state after cloacal vaccination. When PCA was examined separately at each timepoint (**Fig. S1**), group separation emerged by DPI 4 following *H. meleagridis* challenge and was most pronounced at DPI 7-14 in both compartments. PERMANOVA confirmed that treatment was a major driver of metabolic variance in both compartments, with explained variance increasing from 27.4% at DPI 0 to a peak of 58.0% at DPI 14 in cecal tissue, and from 40.2% at DPI 0 to 57.6% at DPI 7 in cecal digesta (**Fig. S2**).

**Fig. 2.**
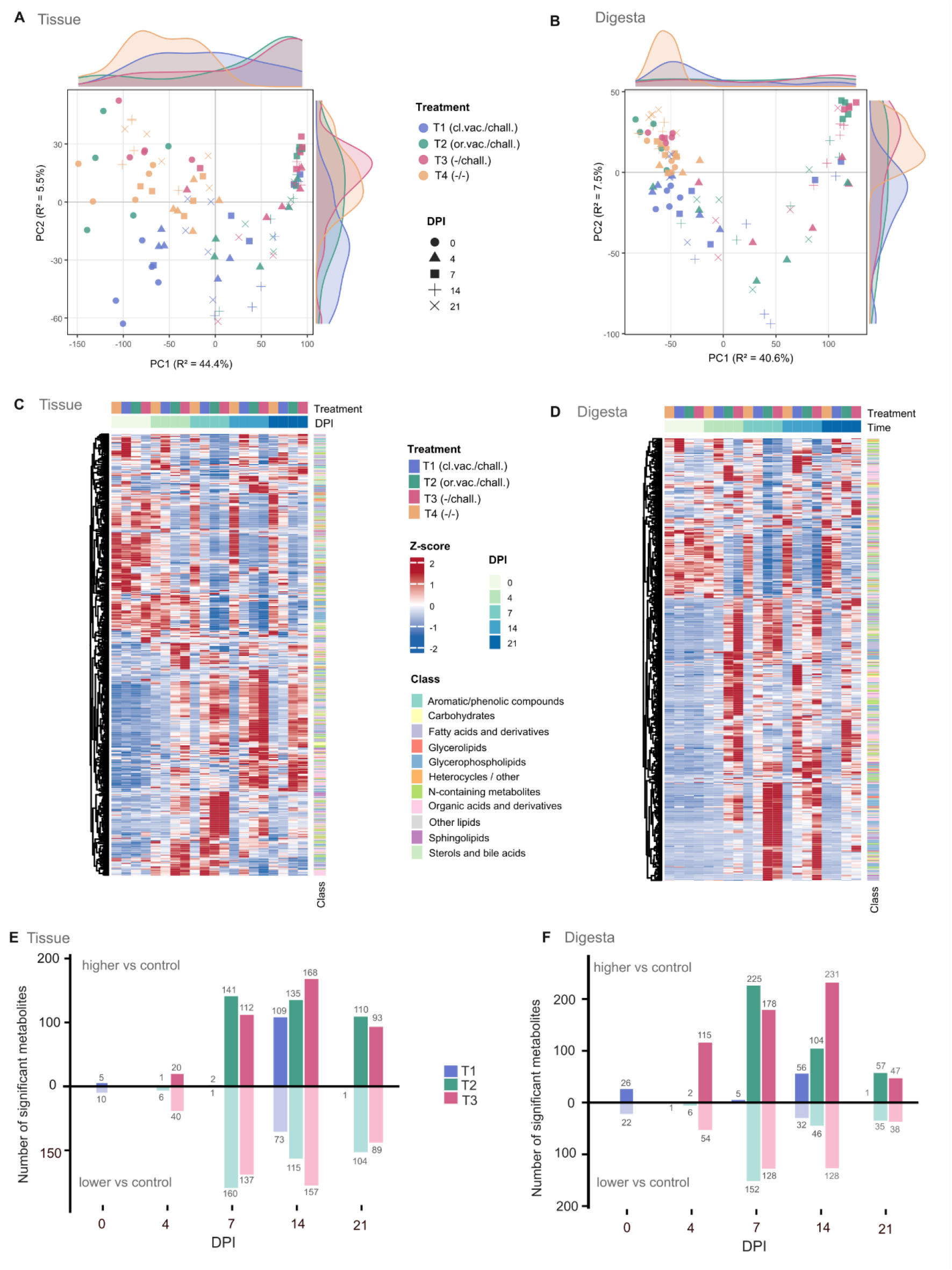
*Histomonas meleagridis* infection drives broad, time-dependent remodeling of chicken cecal metabolome. **(A, B)** Principal component analysis (PCA) of high-quality features in cecal tissue and cecal digesta, respectively, across all treatment groups (T1: cloacal vaccination/challenge; T2: oral vaccination/challenge; T3: -/challenge; T4: -/-) and days post-infection (DPI). A four-component PCA model was retained for both sample types. The tissue model explained 58.7% of the total variance (cumulative R^2^ = 58%, Q^2^ = 54%), while the digesta model explained 56.0% (cumulative R^2^ = 56%, Q^2^ = 51%). PC1 explained 44% and 41% of the total variance in tissue and digesta, respectively. Group separation increased progressively following *H. meleagridis* challenge, becoming most pronounced from DPI 7 onwards. **(C, D)** Unsupervised hierarchical clustering heatmaps of identified metabolites in cecal tissue and cecal digesta, displayed as row-scaled Z scores. Coherent clusters of elevated and suppressed metabolites are visible in infected groups from DPI 4, with vaccinated groups (T1, T2) displaying intermediate profiles relative to T3 and T4. (**E, F**) Number of significantly altered identified metabolites (adj.p < 0.05 vs T4, Benjamini-Hochberg correction) per treatment group across DPI in cecal tissue and digesta.

To examine whether these changes followed broader temporal and treatment-dependent structure, we performed unsupervised hierarchical clustering for all identified metabolites. This analysis revealed a clear treatment- and time-structured disruption in both tissue and digesta (**Fig. 2C-D**). In both compartments, T3 showed the strongest deviation from T4, with broad clusters of metabolites becoming elevated or depleted from DPI 4 onward and generally reaching their greatest divergence at DPI 7 or DPI 14. T1 more often retained a metabolite profile closer to T4, whereas T2 was more frequently intermediate and, in several regions of the heatmap, closer to T3 than to T4. At peak of disease, the disrupted metabolites spanned all major chemical superclasses in both compartments, with organic acids and derivatives, N-containing metabolites, fatty acids and derivatives, and glycerophospholipids most extensively affected (**Fig. S3**). Notably, glycerophospholipids showed a compartment-specific pattern, being predominantly elevated in digesta but more evenly split between elevated and reduced in tissue.

We measured the extent of this disruption by counting significantly altered metabolites relative to the uninfected control (T4; adj.p < 0.05; **Fig. 2E-F**). In challenged unvaccinated birds (T3), the number of altered metabolites increased progressively following infection, peaking at DPI 14 in both cecal tissue and digesta before declining by DPI 21. A substantial number of metabolites were both elevated and reduced, indicating that infection affected metabolism in both directions. Vaccination markedly changed these temporal trajectories. Cloacally vaccinated birds (T1) remained close to T4 through DPI 7 and showed a delayed increase in altered metabolites at DPI 14 that had largely resolved by DPI 21. In contrast, orally vaccinated birds (T2) showed an earlier and broader response, with the number of altered metabolites peaking at DPI 7 and exceeding T3 at that timepoint, before declining thereafter.

### 3.2. *H. meleagridis* infection disrupts microbiota-associated metabolic pathways in the cecum

We next examined metabolites most closely linked to microbial activity in the cecum. Across both tissue and digesta, *H. meleagridis* infection was associated with broad depletion of microbiota-related metabolites, spanning organic acid catabolism, tryptophan metabolism, phytochemical biotransformation, bilirubin reduction, aromatic amino acid metabolism, vitamin-related pathways, and bile acid transformation (**Fig. 3A**). We also observed a smaller set of microbial catabolites and dietary substrates accumulated in infected birds, suggesting that infection did not simply suppress all microbial metabolism but instead reshaped it in a selective manner.

**Fig. 3.**
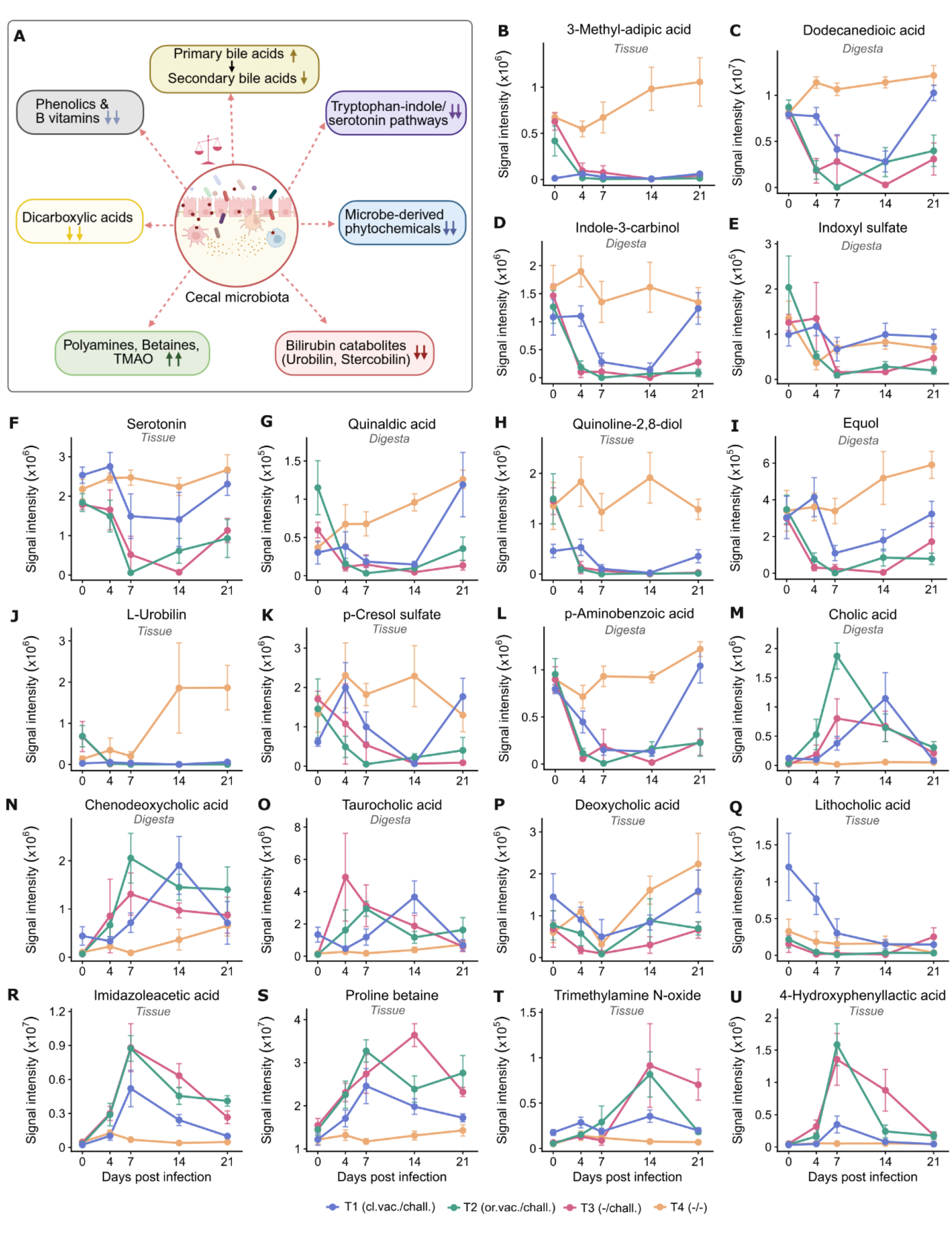
Temporal changes in microbiota-associated metabolites during *H. meleagridis* infection. **(A)** Schematic summary of the major microbiota-associated metabolic processes altered during infection (created with Biorender.com). **(B-U)** Representative temporal profiles of selected microbiota-associated metabolites in cecal tissue or digesta across days post-infection (DPI), including 3-methyl-adipic acid **(B)**, dodecanedioic acid **(C)**, indole-3-carbinol **(D)**, indoxyl sulfate **(E)**, serotonin **(F)**, quinaldic acid **(G)**, quinoline-2,8-diol **(H)**, equol **(I)**, L-urobilin **(J)**, p-cresol sulfate **(K)**, p-aminobenzoic acid **(L)**, cholic acid **(M)**, chenodeoxycholic acid **(N)**, taurocholic acid **(O)**, deoxycholic acid **(P)**, lithocholic acid **(Q)**, imidazolacetic acid **(R)**, proline betaine **(S)**, trimethylamine N-oxide (TMAO) **(T)**, and 4-hydroxyphenyllactic acid **(U)**; full statistical results for displayed metabolites are provided in **Table S2**. Data are presented as mean ± SEM for each treatment group (T1: cloacal vaccination/challenge; T2: oral vaccination/challenge; T3: -/challenge; T4: -/-) at each time point.

One of the clearest patterns was the coordinated loss of dicarboxylic acids and related microbial organic acid catabolites. Eight medium- and long-chain dicarboxylic acids, including adipic, pimelic, suberic, azelaic, sebacic, undecanedioic, dodecanedioic, and 3-methyl-adipic acid, were consistently lower in T3 than in T4 across both cecal compartments (**Fig. 3B-C**; **Fig. S4** and **Table S2**).

Tryptophan metabolism was also broadly disrupted (**Fig. 3D-G**; **Fig. S5** and **Table S2**). Several indole-derived metabolites were depleted in infected birds; for example, indole-3-carbinol (**Fig. 3D**) and indoxyl sulfate (**Fig. 3E**) were significantly lower in both tissue and digesta at DPI 14. Other metabolites, including indoleacetic acid, indoleacetaldehyde, and indole-3-carboxylic acid (**Fig. S5**), were also broadly reduced. Notably, indole-3-propionic acid (**Fig. S5**) remained significantly depleted in tissue from DPI 4 onward, while digesta showed a similar downward trend but did not reach statistical significance. Serotonin (**Fig. 3F**) and N-acetylserotonin were markedly reduced, and quinaldic acid remained low through DPI 21 (**Fig. 3G**). The main exception was indolelactic acid, which increased at DPI 7 in both tissue and digesta (**Fig. S5**).

Of disrupted metabolites, quinoline-2,8-diol, previously reported as a product of microbiota metabolism, though its precise biosynthetic origin within the cecal microbiota remains unclear (Hubbard et al., 2019), showed one of the strongest and most sustained loss in the dataset (**Fig. 3H**). It was already depleted at DPI 4 in both tissue and digesta and remained low throughout the full post-challenge period in the *H. meleagridis* infected groups when compared to the controls.

We also observed a marked depletion of metabolites produced by microbial transformation of dietary compounds (**Fig. 3I**; **Fig. S6** and **Table S2**). For example, equol (**Fig. 3I**), a bacterial metabolite derived from the soy isoflavone daidzein (Iino et al., 2026), was largely depleted from DPI 4 onward in the infected birds. Enterolactone, a microbial product of lignan metabolism (Baldi et al., 2023), showed similarly strong depletion across DPIs. Jasmonic acid, a plant-derived oxylipin catabolized by gut microbiota (Neff and Radka, 2025), was also markedly reduced and remained low through DPI 21.

Microbial bilirubin catabolism was disrupted with similar severity (**Fig. 3J**; **Fig. S6** and **Table S2**). L-urobilin and stercobilin levels were significantly reduced at DPI 14 and 21 in both tissue and digesta. In uninfected control birds, both metabolites increased from DPI 0 to DPI 21, whereas this age-related increase was absent in infected birds (Hall et al., 2024). Instead, they remained depleted throughout the post-challenge period.

Several phenolic and vitamin-related metabolites (e.g., p-Cresol sulfate, gentisic acid, 3-hydroxyphenylacetic acid, 3-hydroxybenzoic acid, p-aminobenzoic acid, 4-pyridoxic acid, biotin, and pyridoxamine) were also consistently depleted in infected birds (**Fig. 3K-L**; **Fig. S6** and **Table S2**) (Aggrey et al., 2019). *H. meleagridis* infection also altered the bile acid profile in a compartment-specific manner (**Fig. 3M-Q, Table S2**). In digesta, both taurine-conjugated and unconjugated primary bile acids accumulated during active infection. For example, cholic acid (**Fig. 3M**) and chenodeoxycholic acid (**Fig. 3N**) were elevated in T3 digesta at DPI 7 and DPI 14, and their conjugated counterparts, taurocholic acid (**Fig. 3O**) and taurochenodeoxycholic acid, showed a similar pattern. By contrast, cholic acid, taurocholic acid, and taurochenodeoxycholic acid showed no significant changes in cecal tissue **(Table S2**). In contrast to the primary bile acids, we observed the opposite pattern of secondary bile acids in tissue: deoxycholic acid (**Fig. 3P)** was reduced at DPI 4 and DPI 14, and lithocholic acid (**Fig. 3Q)** was reduced at DPI 14. In the digesta, there was no significant difference in lithocholic acid between T3 birds and the control, and deoxycholic acid showed only a transient increase at DPI 7 **(Table S2**).

Alongside the broad depletion of microbiota-derived products, infection was associated with the selective accumulation of a distinct set of microbial catabolites and dietary substrates (**Fig. 3R–U**; **Fig. S7** and **Table S2**). Several biogenic amines, such as imidazoleacetic acid (**Fig. 3R**), increased significantly, along with elevated levels of 4-guanidinobutanoic acid, cadaverine, and putrescine (**Fig. S7**) (Formiga et al., 2025). Dietary quaternary ammonium compounds, e.g., proline betaine (**Fig. 3S**), glycine betaine (**Fig. S7**), and TMAO (**Fig. 3T**), accumulated mainly in digesta, especially during peak infection (Buffa et al., 2022; Koistinen et al., 2019). Finally, products of reductive bacterial aromatic amino acid fermentation, e.g., 4-hydroxyphenyllactic acid (**Fig. 3U**) and phenyllactic acid (**Fig. S7)**, were also elevated (Dodd et al., 2017).

### 3.3. Host tissue damage and metabolic reprogramming in the inflamed cecum

In contrast to the broad depletion of microbiota-associated metabolites described above, infected birds (T3) showed accumulation of a distinct set of host-linked metabolites in both cecal tissue and digesta, including itaconic acid, nucleotides, acylcarnitines, ketoleucine, sialic acid, glutathione, and arachidonate-containing lipids (**Fig. 4A**). These changes pointed to immune activation, epithelial injury, mitochondrial dysfunction, structural breakdown of the mucosa, and extensive lipid remodeling in the inflamed cecum.

**Fig. 4.**
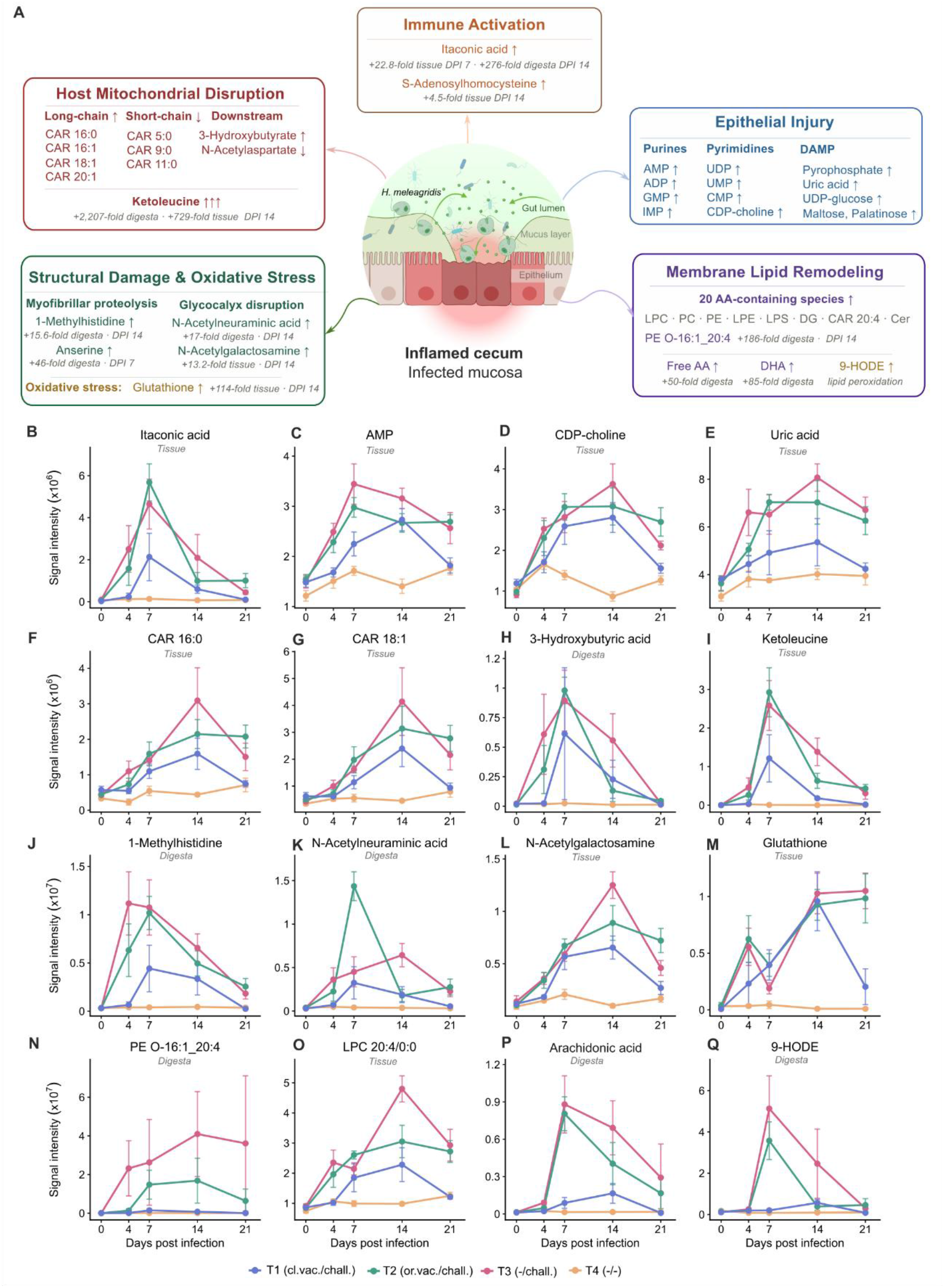
*H. meleagridis* infection is associated with host tissue damage and metabolic reprogramming in the cecum. **(A)** Schematic summary of the major host-associated metabolic responses altered during infection (created with Biorender.com). **(B-Q)** Representative temporal profiles of selected host-associated metabolites in cecal tissue or digesta across days post-infection (DPI), including itaconic acid **(B)**, AMP **(C)**, CDP-choline **(D)**, uric acid **(E)**, CAR 16:0 **(F)**, CAR 18:1 **(G)**, 3-hydroxybutyric acid **(H)**, ketoleucine **(I)**, 1-methylhistidine **(J)**, N-acetylneuraminic acid **(K)**, N-acetylgalactosamine **(L)**, glutathione **(M)**, PE O-16:1_20:4 **(N)**, LPC 20:4/0:0 **(O)**, arachidonic acid **(P)**, and 9-HODE **(Q)**, full statistics for displayed metabolites are provided in **Table S2**. Data are presented as mean ± SEM for each treatment group (T1: cloacal vaccination/challenge; T2: oral vaccination/challenge; T3: -/challenge; T4: -/-) at each timepoint. AA: Arachidonic acid; DHA: Docosahexaenoic acid; LPS: Lysophosphatidylserine; LPC: Lysophosphatidylcholine; PE: Phosphatidylethanolamine; PC: Phosphatidylcholine; CAR: Acylcarnitine; DG: Diacylglycerol; Cer: Ceramide.

Metabolites linked to inflammation and macrophage activation were among the earliest host-associated signals. Itaconic acid (**Fig. 4B)** and S-adenosylhomocysteine (**Fig. S8**) were elevated in infected birds from DPI 4 onward in digesta and from DPI 7 in tissue (**Table S2**).

A broad accumulation of nucleotides, nucleotide sugars, and purine breakdown products was also observed (**Fig. 4C-E**; **Fig. S8** and **Table S2**). Notably, this included several metabolite-derived damage-associated molecular patterns (DAMPs), such as uric acid, succinate, pyrophosphate, and UDP-glucose, alongside other key intermediates, including AMP, CDP-choline, GMP, UDP-glucuronic acid, CMP, and UMP.

The infected cecum also showed a clear signature of altered acylcarnitine metabolism and amino acid catabolism. Long-chain acylcarnitines, including CAR 16:0 (**Fig. 4F**) and CAR 18:1 (**Fig. 4G**), were elevated, while short- and medium-chain acylcarnitines, e.g., CAR 5:0 and CAR 9:0, were reduced (**Fig. S9**). 3-Hydroxybutyric acid (**Fig. 4H**) and ketoleucine (**Fig. 4I**) were both markedly increased in tissue and digesta (**Table S2**). A coordinated depletion of N-acetylated amino acids, including N-acetylglutamic acid, N-acetylhistidine, N-acetylarginine, and N-acetylmethionine, was also observed at DPI 14 (**Fig. S10**).

Several metabolites indicated progressive structural damage to the cecal surface. 1-methylhistidine (**Fig. 4J**), sialic acid (N-acetylneuraminic acid, **Fig. 4K**), N-acetylgalactosamine (**Fig. 4L**), and anserine (**Fig. S10**) were all elevated in infected birds across multiple post-challenge timepoints. Glutathione (**Fig. 4M**) was also markedly increased, particularly in the tissue.

The clearest lipid remodeling signal was PE O-16:1_20:4 (**Fig. 4N, Table S2**), which was elevated at all four post-challenge timepoints in both compartments. More broadly, 20 identified metabolites carrying a 20:4 acyl chain were elevated in T3 relative to T4, spanning lysophosphatidylcholines, e.g., LPC 20:4/0:0 (**Fig. 4O**), phosphatidylcholines, phosphatidylethanolamines, diacylglycerols, acylcarnitines, and ceramides (**Fig. S11**). Free arachidonic acid (FA 20:4, **Fig. 4P**), docosahexaenoic acid (FA 22:6, **Fig. S10**), and 9-HODE (FA 18:2;O, **Fig. 4Q**) were also elevated in both tissue and digesta (**Table S2**).

### 3.4. Protection by vaccination preserves key microbial and host metabolic functions

We then examined how vaccination specifically changed the metabolic response to infection. Cloacal vaccination clearly reduced many of the changes seen in challenged-only birds, and this effect was generally stronger than in the oral vaccination group (**Fig. 5**). In most cases, the protection was more evident in digesta than in the tissue.

**Fig. 5.**
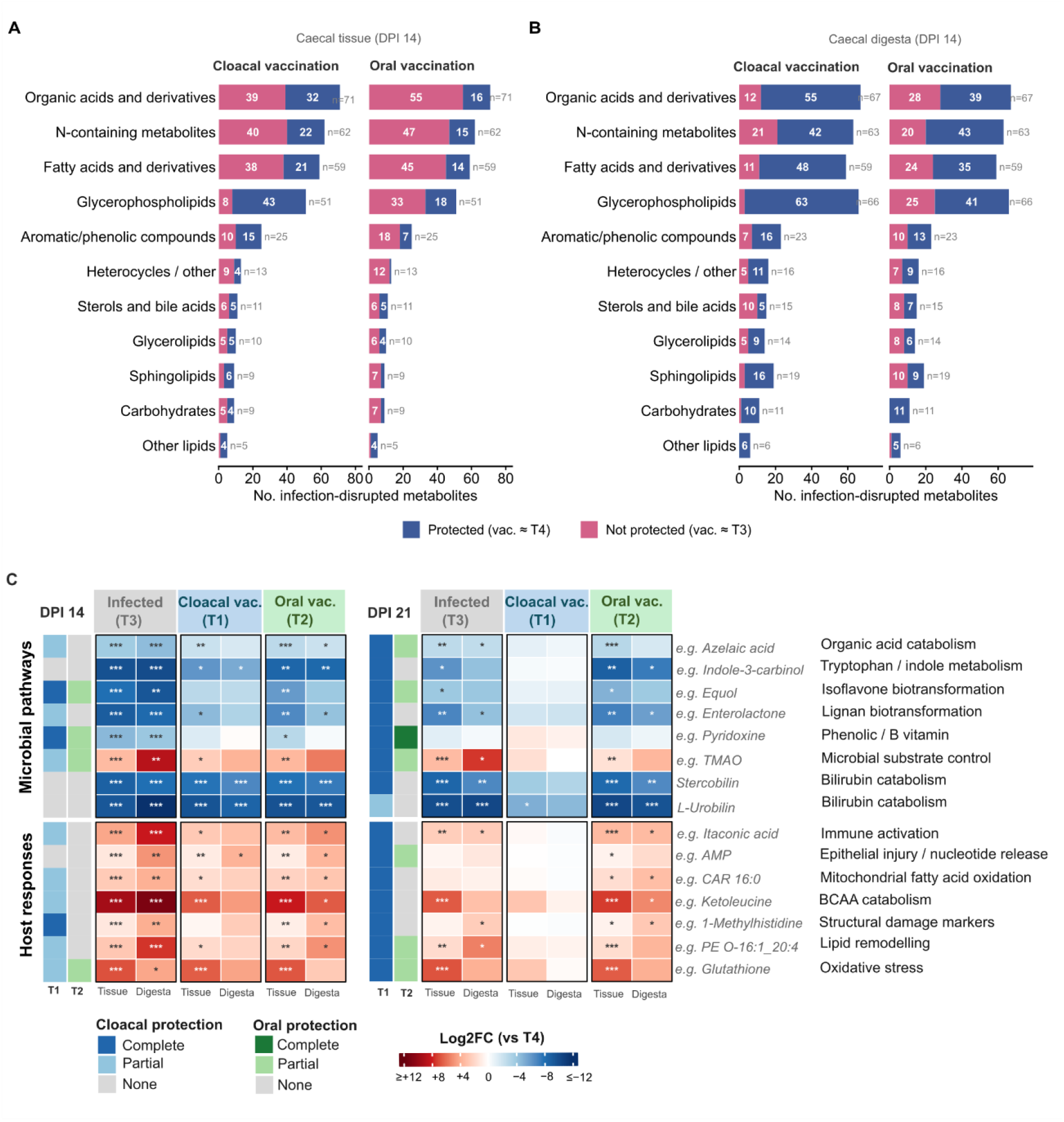
Cecal metabolic protection conferred by cloacal and oral vaccination against *H. meleagridis* challenge. **(A-B)** Protection analysis for infection-disrupted metabolites in cecal tissue **(A)** and cecal digesta **(B)** at DPI 14. Metabolites significantly altered by infection (T3 vs T4, adj.p < 0.05) were classified as protected by vaccination route (vaccinated group not significantly different from T4, adj.p ≥ 0.05; blue) or not protected (vaccinated group significantly altered in the same direction as T3, adj.p < 0.05; pink). Bars show the number of disrupted metabolites per chemical superclass (n indicates total per class). **(C)** Heatmaps showing log_2_ fold changes (log_2_FC) of representative metabolites from pathway modules in vaccinated challenged groups (T1, cloacal; T2, oral) and the unvaccinated challenged group (T3), relative to the negative control (T4), at peak infection (DPI 14) and during recovery (DPI 21). Each row corresponds to one pathway module, with the representative metabolite indicated in italics (one per module; full statistics in **Table S2**). Color intensity reflects log_2_FC from depletion (blue, ≤ −12) to elevation (red, ≥ 12), using the same scale across panels. Left color bars summarize protection status for T1 (blue) and T2 (green) at each timepoint as Complete (no significant change in either compartment), Partial (significant in one compartment), or None (significant in both tissue and digesta). Significance: * adj.p < 0.05, ** adj.p < 0.01, *** adj.p < 0.001.

Before *H. meleagridis* challenge (DPI 0), cloacally vaccinated birds (T1) exhibited a distinct metabolic profile, characterized by elevated levels of biliverdin, TMAO, pipecolic acid, and bile acids, e.g., taurochenodeoxycholic acid and lithocholic acid, together with a marked reduction in dicarboxylic acids and long-chain fatty acids, e.g., 3-methyl-adipic acid and elaidic acid (**Fig. S12**). In contrast, orally vaccinated birds (T2) and the unvaccinated challenged group (T3) showed no significant differences from T4 at DPI 0 in either compartment, indicating that the pre-challenge metabolic shift was specific to the cloacal vaccination route.

To estimate the extent of vaccine-mediated metabolic protection at peak infection, we classified infection-disrupted metabolites as protected if they were no longer significantly altered in the vaccinated group relative to T4 (adj.p ≥ 0.05; **Fig. 5A-B**). The results showed that cloacal vaccination (T1, 49% in tissue, and 78% in digesta) conferred broader protection against infection-disrupted identified metabolites than oral vaccination (T2, 27% in tissue, and 61% in digesta). Notably, the clearest route-dependent difference was seen for lipid-related classes.

Cloacal vaccination preserved several microbial metabolic functions that were disrupted in T3 (**Fig. 5C**, **Table S2**). Among microbiota-related metabolites, equol and enterolactone were no longer significantly different from T4 in T1 digesta, while jasmonic acid showed a similar pattern. Tryptophan-related metabolites, such as serotonin and indole-3-carbinol, were also better preserved in T1. Several metabolites that accumulated in T3 digesta, e.g., TMAO, glycine betaine, cadaverine, and 4-guanidinobutanoic acid, were no longer significantly elevated.

However, we observed that protection was not complete. Quinoline-2,8-diol (**Fig. 3H**) remained profoundly depleted in T1, and both stercobilin and L-urobilin also remained strongly reduced at DPI 14 (**Fig. 5C**, **Table S2**). This was especially clear for bilirubin-related metabolites, which were poorly restored in both vaccinated groups. Bile acid disturbances were also only partly corrected (**Fig. 3M-Q)**.

Host-associated changes were also attenuated by cloacal vaccination (**Fig. 4B-Q**, **Fig. 5C, Table S2**). For example, itaconic acid was no longer significantly elevated in T1 digesta at DPI 14, and ketoleucine was markedly reduced relative to T3, though it remained significantly elevated in tissue. Markers of structural damage and epithelial injury, e.g., 1-methylhistidine and AMP, showed partial to near-complete protection in T1. Lipid remodeling markers were especially well controlled in T1 digesta, where PE O-16:1_20:4 and LPC 20:4/0:0 were no longer significantly elevated. Protection in tissue, however, was more partial. Glutathione was the clearest exception, remaining strongly elevated in both T1 and T3 tissue.

Oral vaccination showed the same overall direction of protection, but the effect was consistently weaker, most evidently for host-associated metabolites, where, e.g., itaconic acid, ketoleucine, CAR 16:0, and PE O-16:1_20:4 remained significantly elevated in T2 digesta at DPI 14.

By DPI 21, the difference between vaccination routes was pronounced (**Fig. 5C**). In cloacally vaccinated birds, the cecal metabolome returned close to the control state across several metabolic pathways. Microbiota-related metabolites, including equol, indole-3-carbinol, and pyridoxine, and host-associated markers, e.g., itaconic acid, AMP, ketoleucine, 1-methylhistidine, PE O-16:1_20:4, and glutathione, were no longer significantly different from T4. Meanwhile, L-urobilin was still depleted in tissue at DPI 21. In contrast, orally vaccinated birds showed a more persistent and incomplete metabolic recovery than those vaccinated by the cloacal route. Metabolic disruption remained evident at DPI 21 across multiple pathway modules in both tissue and digesta, including tryptophan and indole metabolism, immune activation, acylcarnitine profiles, structural damage markers, and bilirubin catabolites.

## 4. Discussion

Histomonosis has long been described through pathology, parasitology, and even microbiology, but its local metabolic consequences in the cecum have been poorly defined (Lesleigh C. Beer et al., 2022; Dubey et al., 2024; Chen et al., 2026). Here we show broad disruption of both microbiota and host-related metabolite profiles in the cecum. By analyzing both cecal tissue and digesta across challenge with or without vaccination, and recovery, this study provides a spatially and temporally resolved metabolic view of histomonosis. The main finding is that *H. meleagridis* infection disrupted the cecal host-microbiota metabolic interface by broad loss of microbiota-associated metabolites, accompanied by a host response consistent with mucosal injury and inflammatory metabolic stress. These changes were strongest during DPI 7-14, and especially, cloacal vaccination was able to substantially reduce the *H. meleagridis* infection-caused metabolic disruption.

Previous metabolomics work on *H. meleagridis* was limited to parasite culture or systemic plasma/liver analysis in a mixed-infection model, and therefore did not resolve the infected cecal microenvironment itself (Ammar et al., 2024; Oladosu et al., 2023). Despite these very different experimental settings, several findings were consistent with our analysis, including altered TMAO, succinate, and tryptophan-related metabolism. However, direct profiling of cecal tissue and luminal digesta revealed substantially broader disruption of microbiota-associated metabolism, including marked depletion of indole metabolites, bilirubin catabolites, bile acid transformation products, and microbial organic acid pathways that had not been resolved before.

Microbiome studies have established that *H. meleagridis* infection disrupts the cecal microbial community in both chickens and turkeys. Specifically, in chickens, *H. meleagridis* infection has been associated with reduced microbial diversity, clear separation of infected and control cecal microbiota, depletion of commensal taxa such as Lactobacillus-related species and *Faecalibacterium prausnitzii*, and enrichment of dysbiosis-associated taxa including Proteobacteria, *Escherichia*, *Bacteroides*, and *Fusobacterium*, most evident around DPI 14 (Abdelhamid et al., 2020; Chen et al., 2026). Our metabolomics data extend those observations by linking structural dysbiosis to a broad functional reorganization of microbial metabolism. The coordinated depletion of medium- and long-chain dicarboxylic acids, such as 3-methyl-adipic acid and dodecanedioic acid, points to perturbations in fatty acid ω-oxidation and related organic acid catabolism, which may reflect disrupted host-microbiota cross-talk in lipid metabolic pathways (Franzosa et al., 2019). The broad disruption of tryptophan-derived metabolites spanning the indole, serotonin, and kynurenine branches, together with depletion of equol, enterolactone, and bilirubin catabolites, further reflects a loss of functionally diverse microbial transforming activity, potentially compounded by reduced feed intake during infection (Agus et al., 2018; Baldi et al., 2023; Hall et al., 2024; Iino et al., 2026; Kipp et al., 2025; Sinha et al., 2024; Zhang et al., 2026). The compartment-specific bile acid pattern, primary bile acid accumulation in digesta alongside secondary bile acid depletion in tissue, is consistent with disturbed microbial bile acid transformation, including pathways associated with deconjugation and 7α-dehydroxylation, and may additionally reflect disturbed epithelial bile acid reabsorption secondary to mucosal damage (Bansal et al., 2020; Zhang et al., 2025).

At the same time, the selective accumulation of several metabolites, e.g., imidazoleacetic acid, cadaverine, putrescine, betaines, and TMAO, indicates that infection did not uniformly silence microbial activity but disrupted host-microbe interaction, redirecting metabolism toward specific catabolic routes. Cadaverine and putrescine have previously been reported as products of microbial amino acid decarboxylation, but are also associated with tissue decomposition; their accumulation may partly reflect microbial access to host-derived substrates through a damaged mucosal layer (Franzosa et al., 2019). The accumulation of proline betaine, glycine betaine, and TMAO, compounds normally processed by gut bacteria, further supports the selective reorganization of microbial quaternary amine metabolism (Buffa et al., 2022; Koistinen et al., 2019). Together, these findings indicate that *H. meleagridis* infection did not simply suppress microbial metabolism but reorganized it in a pathway-specific manner, consistent with the broader cecal dysbiosis reported in previous microbiome studies.

The host-associated metabolite pattern closely matched the clinical and pathological course observed in the trial (Hatfaludi et al., in prep.). Clinical and pathological findings showed that challenged-only birds had reduced body weight, more pronounced clinical signs from DPI 7 onward, and increasing cecal lesion severity during the first week after challenge, whereas vaccination reduced disease burden, especially after cloacal delivery, pointing towards importance of vaccine delivery. Consequently, this study provides a biochemical view of this host response. The early rise in itaconic acid suggests that innate immune activation was already present before peak cecal pathology, consistent with the established role of ACOD1/IRG1-derived itaconate in macrophage metabolic reprogramming during inflammation (Lampropoulou et al., 2016). The later accumulation of nucleotides, nucleotide sugars, uric acid, and pyrophosphate in digesta is consistent with epithelial injury and release of intracellular metabolites into the lumen (Kang et al., 2024; Kono and Rock, 2008). In parallel, long-chain acylcarnitines, ketoleucine, 3-hydroxybutyric acid, and depleted N-acetylated amino acids point to disturbed mitochondrial and amino acid metabolism in the inflamed cecum, consistent with broader evidence linking intestinal inflammation to altered mitochondrial metabolism (Biswas et al., 2020; Joshi et al., 2006; Newman and Verdin, 2017; Pessentheiner et al., 2013; Tang et al., 2025). Structural and oxidative injury were further supported by increased sialic acid, a marker of mucosal glycoprotein disruption (Arias et al., 2025; Linden et al., 2008), elevated 1-methylhistidine and anserine, indicative of breakdown of methylhistidine-containing structural proteins and avian muscle-associated dipeptides (Barbaresi et al., 2019; Sjölin et al., 1987), and increased glutathione, consistent with a strong oxidative stress response in the inflamed mucosa (Franco and Cidlowski, 2009; Perricone et al., 2009). The coordinated elevation of arachidonate-containing lipids across multiple lipid classes, together with free arachidonic acid, DHA, and oxidized lipids such as 9-HODE, indicates extensive membrane phospholipid remodeling and oxidative lipid damage during active infection (Das, 2018; Gilroy and Bishop-Bailey, 2019; Spiteller and Spiteller, 1997).

Vaccination modified both the magnitude and the duration of the cecal metabolic disturbance. Evidence for vaccination against histomonosis in chickens remains limited. One study demonstrated vaccination of pullets prior to lay with *in vitro* attenuated *H. meleagridis* five weeks before challenge reduced the decline in egg production and lowered the incidence and severity of pathological lesions, although protection was incomplete comparing with non-challenged birds (Liebhart et al., 2013). Existing immunological work suggests that protection involves controlled local responses in the cecum, including IFN-γ-associated cellular and TLR-mediated innate responses, but has not defined how vaccination reshapes the biochemical environment of cecal tissue and lumen (Kidane et al., 2018; Mitra et al., 2021, 2018). Our results show that this protection is clearly visible at the metabolic level. At peak of disease, cloacal vaccination (T1) protected a larger fraction of infection-disrupted metabolites than oral vaccination (T2). In digesta, 78% of disrupted metabolites were protected by cloacal vaccination compared with 61% in orally vaccinated birds, and the same overall pattern was also evident in cecal tissue samples. This difference was most apparent for host-associated damage markers and lipid-related metabolites, effects which were much less altered after cloacal vaccination but often remained significantly different after oral vaccination. The recovery phase further separated the two routes. By DPI 21, most microbiota- and host-associated markers in cloacal vaccination group had returned to a state close to the control, whereas oral vaccination group retained broader metabolic disturbances across tryptophan metabolism, immune activation, mitochondrial and amino acid metabolism, structural damage markers, and bilirubin catabolites. This was in line with clinical observations, in which cloacally vaccinated birds had lower clinical scores and complete resolution of cecal lesions by three weeks post-challenge, whereas residual lesions remained detectable after oral vaccination.

An additional finding was that cloacally vaccinated birds already differed metabolically from controls before virulent challenge. As DPI 0 samples were collected after vaccination, this represents a post-vaccination, pre-challenge state rather than a true baseline. The involvement of several microbiota-associated metabolites suggests that administration of attenuated *H. meleagridis* may have modified the cecal biochemical environment before challenge. However, without pre-vaccination samples or concurrent microbiome and immune profiling, it remains unclear whether this shift reflected microbial restructuring, local immune priming, exposure to the attenuated parasite, or a combination of these processes. Moreover, the persistence of selected disturbances, particularly bilirubin and bile acid metabolism, indicates that cloacal vaccination reduced disease-associated metabolic disruption without fully restoring all cecal functions.

The strengths of our study include its longitudinal vaccination-challenge design, which enabled the temporal development and recovery of metabolic responses to be tracked during infection, and the parallel profiling of cecal tissue and digesta, which provided complementary information on host- and microbiota-associated metabolic changes. The inclusion of two vaccination routes further allowed us to compare differences in the magnitude and timing of metabolic protection within the same experimental framework. Nevertheless, some limitations should be noted in this study. One limitation of the study is the modest number of birds analyzed at each time point. Although this was sufficient to detect clear and consistent pathway-level changes, it may reduce confidence in interpreting individual metabolites with smaller or more variable effects. In addition, future study designs that include sampling before vaccination and during the interval between vaccination and challenge may provide further insight into metabolic changes induced by vaccination prior to challenge. Also, the mechanisms underlying these metabolite changes cannot be resolved by metabolomics alone and thus will require future investigation with microbiome and targeted immunological analyses. Extending this framework to turkeys in future studies is also essential, given the greater severity of histomonosis in turkeys, to define which aspects of the metabolic response to histomonosis are conserved and which are host-specific.

## 5. Conclusion

This study shows that *H. meleagridis* infection causes a coordinated disruption of the cecal host-microbiota metabolic interface, characterized by a broad loss of microbiota-associated metabolic functions and a distinct host injury response. By profiling both cecal tissue and luminal digesta during infection and vaccination, we show that cloacal vaccination attenuated this disruption more effectively than oral vaccination and was associated with faster metabolic recovery. These findings provide the first cecum-focused metabolomic map of *H. meleagridis* infection and vaccination in chickens, offering insight into host-microbe interactions and the role of vaccination at the metabolic scale. More broadly, our findings suggest that effective control of histomonosis depends not only on limiting parasite-induced tissue damage but also on preserving or restoring key microbial functions in the cecum, which may inform future vaccine refinement and complementary microbiota-targeted strategies in poultry production.

## 6. Declarations

### 6.1. Humane care of animals/animal ethics

The animal trial and all the included procedures on experimental birds were discussed and approved by the institutional ethics committee of the University of Veterinary Medicine, Vienna and licensed by the Austrian Federal Ministry of Education, Science (ref: BMBWF 2024-0.062.643)

### 6.2. Data Availability

The metabolomics datasets generated and analyzed during this study are publicly available in the MetaboLights repository under accession number **MTBLS13488** (https://www.ebi.ac.uk/metabolights/MTBLS13488), while further details and data from the animal experiment can be found at www.3domics.eu/database (Lee et al., under review).

### 6.3. Supplementary Material

Supplementary information is available with this manuscript and includes the following files:

- **Supplementary File 1**: **Supplementary Methods** detailing sample preparation, LC–MS data acquisition, and data preprocessing workflows; **Supplementary Table S1** summarizing the number of high-quality metabolomic features detected across analytical modes and sample types; and **Supplementary Figures S1-S12** providing additional multivariate analyses, statistical summaries, and extended temporal metabolite profiles across tissues, digesta, treatment groups, and timepoints.
- **Supplementary File 2**: **Supplementary Table S2** containing the complete metabolite identification results and corresponding statistical analyses for identified metabolites, including hierarchical chemical classification, database identifiers, log2 fold changes, and adjusted p-values.

### 6.6. CRediT authorship contribution statement

**Anh Vu Nguyen**: Formal analysis, Data curation, Investigation (metabolite identification and curation), Visualization, Writing – original draft. **Topi Meuronen**: Data curation, Formal analysis, Investigation (LC-MS analysis, metabolite identification and curation), Writing – review & editing. **Atte Lihtamo**: Formal analysis, Data curation, Writing – review & editing. **Soile Turunen**: Investigation (LC-MS analysis), Writing – review & editing. **Tamas Hatfaludi**: Conceptualization, Investigation (animal experiment), Resources, Ethics approval and regulatory compliance, Writing – review & editing. **Michael Hess**: Conceptualization, Investigation (animal experiment), Resources, Ethics approval and regulatory compliance, Funding acquisition, Writing – review & editing. **Antton Alberdi**: Conceptualization, Funding acquisition, Project administration, Writing – review & editing. **Kati Hanhineva**: Conceptualization, Supervision, Funding acquisition, Writing – review & editing. **Olli Kärkkäinen**: Conceptualization, Supervision, Project administration, Funding acquisition, Writing – review & editing.

## 6.7. Acknowledgments

The authors thank the staff of the animal facilities at the University of Veterinary Medicine Vienna for their assistance with animal care, sample collection, and technical support during the experimental trial. The authors also thank Dr Amalia Bogri and Dr Carlotta Pietroni at the University of Copenhagen for their assistance in processing and preparation of samples for metabolomics analysis. The authors further acknowledge all contributors to the 3D-omics and HoloGen projects for valuable discussions and support.

## 6.8. Declaration of generative AI and AI-assisted technologies in the manuscript preparation process

During the preparation of this manuscript, the authors used ChatGPT to assist with language polishing, clarity, and stylistic improvements. Grammarly was also used for grammar and spelling checks. After using these tools, the authors thoroughly reviewed and edited the content and take full responsibility for the content of the published article.

