## Supplementary file 1 for "*Histomonas meleagridis* infection and vaccination protection in chickens – a metabolomics study"

Assoc. Professor Olli Kärkkäinen,

School of Pharmacy, Faculty of Health Sciences, University of Eastern Finland,

70210 Kuopio, Finland

### Supplementary Methods

*Details of Sample preparation, LC-MS parameters and data preprocessing*

##### *Sample Preparation*

Frozen cecal tissue and digesta samples were weighed into pre-cooled bead homogenizer tubes (Bead Ruptor Pre-Filled Bead Tubes, 2 mL, metal beads; Omni International). Metabolites were extracted by solid-liquid extraction using 80% methanol (v/v; CHROMASOLV LC-MS Ultra, Riedel-de Haën, Honeywell) in ultrapure water (Type 1; Direct-Q Water Purification System) at standardized sample-to-solvent ratios: 1000 µL per 100 mg tissue and 1500 µL per 100 mg digesta. Extraction blanks were prepared identically using 1600 µL of solvent per tube. Tissue samples were homogenized at 0-4°C using a bead mill homogenizer (Bead Ruptor 24 Elite with cryo unit; Omni International) at 7.45 m/s for two consecutive 60-second cycles; digesta samples were processed for a single cycle. Extracts were incubated on ice for 15 minutes with one vortex mixing step (10 seconds), then centrifuged at 17,000 × g for 10 minutes at 4°C. Supernatants were transferred to a 0.2 µm filter plate (Captiva ND, Agilent) and centrifuged at 700 × g for 5 minutes at 4 ºC to remove particulates. Pooled QC samples were prepared separately for each sample type by combining 50 µL aliquots from all supernatants of the corresponding type, filtered identically, and used throughout for instrument conditioning, drift correction, and quality monitoring. Prepared extracts were stored at 2-8°C and analyzed on the same day or within a short period.

##### *LC-MS data acquisition*

LC-MS analysis was performed on an Agilent 1290 Infinity II UHPLC coupled to an Agilent 6546 QTOF mass spectrometer with a Dual Jet Stream electrospray ionization (ESI) source (REF). To maximize metabolome coverage, each sample was analyzed in four complementary analytical modes: reversed-phase (RP) chromatography and hydrophilic interaction liquid chromatography (HILIC) in positive and negative ionization modes.

RP separation was performed on a Zorbax Eclipse XDB-C18 column (2.1 × 100 mm, 1.8 µm; Agilent Technologies) at 50°C with a flow rate of 0.4 mL/min over a 16.5-minute gradient using water (eluent A) and methanol (eluent B), both with 0.1% v/v formic acid. The gradient program was as follows (time [min], %B): (0, 2), (10, 100), (14.5, 100), (14.51, 2), (16.5, 2). HILIC separation was performed on an Acquity UPLC BEH Amide column (2.1 × 100 mm, 1.7 µm; Waters Corporation) at 45°C with a flow rate of 0.6 mL/min over a 12.5-minute gradient using 50% (v/v) acetonitrile in water with 20 mM ammonium formate and 0.25% v/v formic acid (eluent A) and 90% (v/v) acetonitrile in water with 20 mM ammonium formate and 0.25% v/v formic acid (eluent B). The gradient program was as follows (time [min], %B): (0, 100), (2.5, 100), (10, 0), (10.01, 100), (12.5, 100). For RP separations, the initial 1.0 min was diverted to waste; for HILIC separations, data acquisition began at 0.0 min.

ESI source parameters were identical across all analytical modes: drying gas temperature 325°C, drying gas flow 10 L/min, nebulizer pressure 45 psig, sheath gas temperature 350°C, sheath gas flow 11 L/min, capillary voltage (VCap) 3500 V, nozzle voltage 1000 V, fragmentor 100 V, skimmer 45 V, and octopole RF peak 750 V.

Full MS data were acquired across a scan range of 20-1600 m/z at a scan rate of 1.67 spectra/sec. Tandem MS (MS/MS) data were acquired by data-dependent acquisition (DDA) at a scan rate of 3.33 spectra/sec, selecting up to four precursor ions per cycle for fragmentation at collision energies of 10, 20, and 40 eV, with an isolation width of approximately 1.3 amu. Precursors were sorted by abundance and selected above an absolute threshold of 200 counts. Each precursor was excluded from reselection after two MS/MS acquisitions and released after 0.25 minutes. Purity cutoff was set to 30% with 100% purity stringency, and isotope modelling used the common isotope model. MS/MS acquisition spanned three static mass exclusion windows (50-249, 250-750, and 601-900 m/z) to improve fragment coverage across the mass range. All raw data were collected in centroid form using MassHunter Acquisition B.10.1 (Agilent Technologies).

Continuous mass axis calibration was maintained using reference ions at m/z 121.050873 and 922.009798 (ESI+) and m/z 119.03632 and 966.000725 (ESI−). Analytical quality was monitored throughout each run by injection of extraction blanks, an LC-MS QC Reference Standard (9-compound mix; Waters Corporation) at the start of each run, and pooled QC samples injected at regular intervals.

***Peak detection and spectral processing***

Raw data were processed in MS-DIAL (v4.9). Peak detection used a linear weighted moving average algorithm with a minimum peak height of 2000 amplitude units, mass slice width of 3000 Da, smoothing level of 3 scans, and minimum peak width of 5 scans. MS1 and MS2 mass tolerances of 0.01 Da and 0.025 Da were applied for data collection. Peak alignment used an MS1 tolerance of 0.015 Da and a retention time tolerance of 0.1 minutes. Blank filtering was not applied during alignment but was performed downstream in R, allowing retention of biological signals co-eluting with background ions.

***Data preprocessing***

Raw peak intensity matrices were imported into R and preprocessed using the notame package. Preprocessing was applied independently to each combination of analytical mode and sample type (cecal tissue, cecal digesta), yielding eight parallel preprocessing streams. Features were excluded if their maximum biological sample intensity was less than five-fold above the mean blank intensity, or if detected in fewer than 70% of QC samples and fewer than 50% of samples within every treatment group. Instrument signal drift was corrected per feature by fitting a smoothed cubic spline to log-transformed QC intensities as a function of injection order; the smoothing parameter was optimized by generalized cross-validation (range 0.5-1.5), and drift correction was applied only where both RSD and D-ratio improved post-correction. Feature quality was assessed using robust QC-derived metrics: features were excluded if the robust relative standard deviation (RSD* = 1.486 × MAD(QC) / median(QC)) exceeded 0.20 or if the dispersion ratio (D-ratio* = MAD(QC) / MAD(biological)) exceeded 0.40. Between-sample normalization used probabilistic quotient normalization (PQN) with the median QC sample as reference. Residual missing values, attributed to intensities below the detection threshold, were replaced with zero. All samples were measured within a single analytical batch.

### Supplementary Tables

**Table S1.** Number of high-quality features* detected across four LC-MS modes.

| Analytical mode | Tissue | Digesta | Detected at least one Tissue/Digesta | Detected in both Tissue&Digesta |
|---|---|---|---|---|
| HILIC_neg | 6093 | 5290 | 7623 | 3760 |
| HILIC_pos | 6268 | 6637 | 8740 | 4165 |
| RP_neg | 6507 | 6975 | 8782 | 4700 |
| RP_pos | 7033 | 8658 | 10699 | 4992 |
| Total | 25901 | 27560 | 35844 | 17617 |

**features that passed all preprocessing quality filters (contaminant removal, detection frequency, drift correction, signal quality) within each analytical mode and sample type. Modes: HILIC = hydrophilic interaction liquid chromatography; RP = reversed-phase; neg/pos = ionization polarity.*

### Supplementary Figures

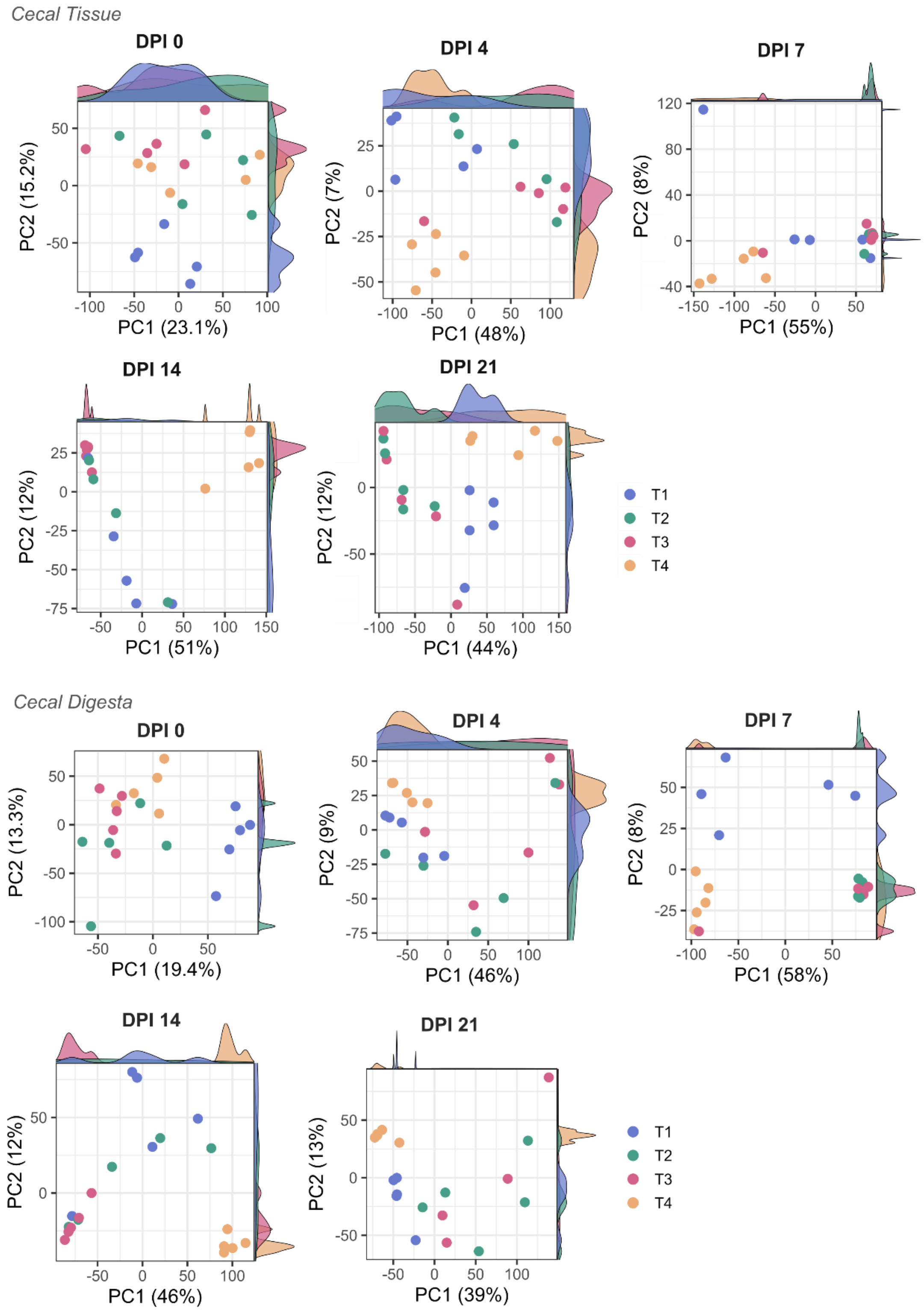


**Fig. S1. Principal components at five DPI: 0, 4, 7, 14, 21 for cecal tissue (upper panels) and digesta (lower panels) across all treatment groups and timepoints**. Each point represents one biological sample, colored by treatment group (T1: cloacal vaccination/challenge; T2: oral vaccination/challenge; T3: challenge only, unvaccinated; T4: negative control, unvaccinated and unchallenged).

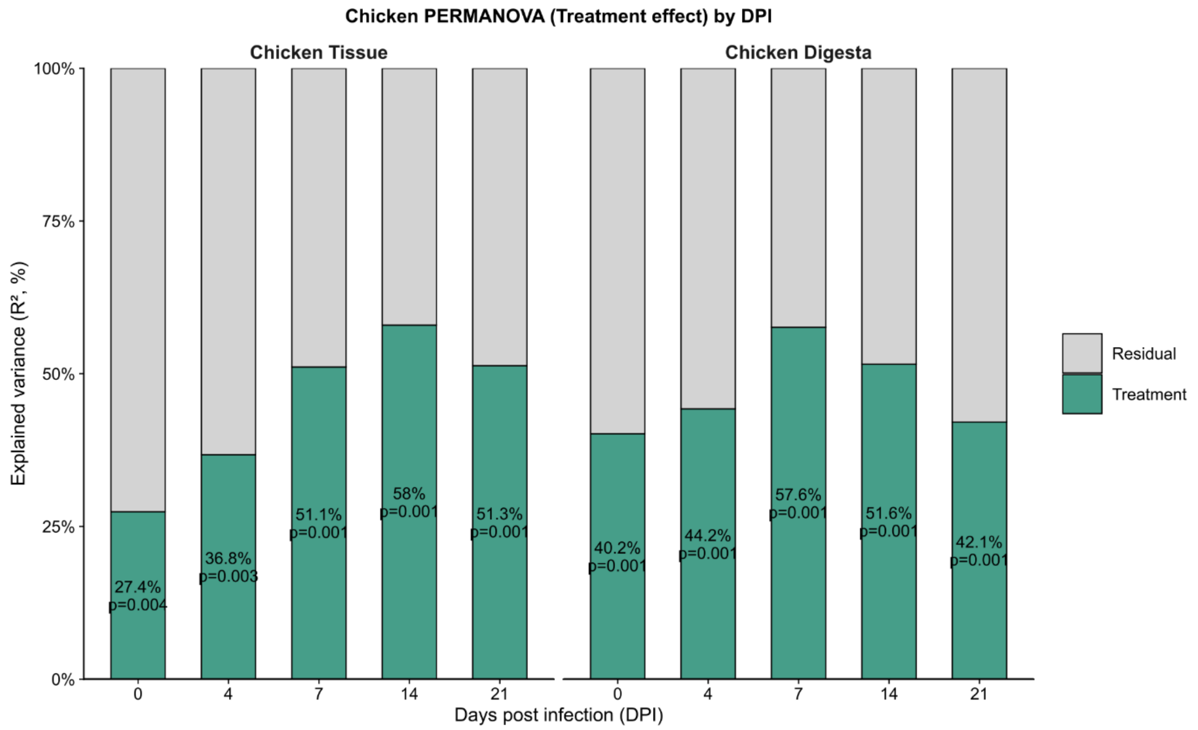


**Fig. S2. PERMANOVA of cecal metabolome profiles across treatments and days post-infection**. Stacked bars show the proportion of variance explained by treatment (T1-T4) versus residual variance at each sampling time point for cecal tissue (left) and digesta (right). Values above blue bars indicate the PERMANOVA $R^2$ and permutation-based p-values, demonstrating that vaccination route and *H. meleagridis* challenge together explain a substantial and statistically significant fraction of metabolome variation, with strongest effects at 7-14 days post infection.

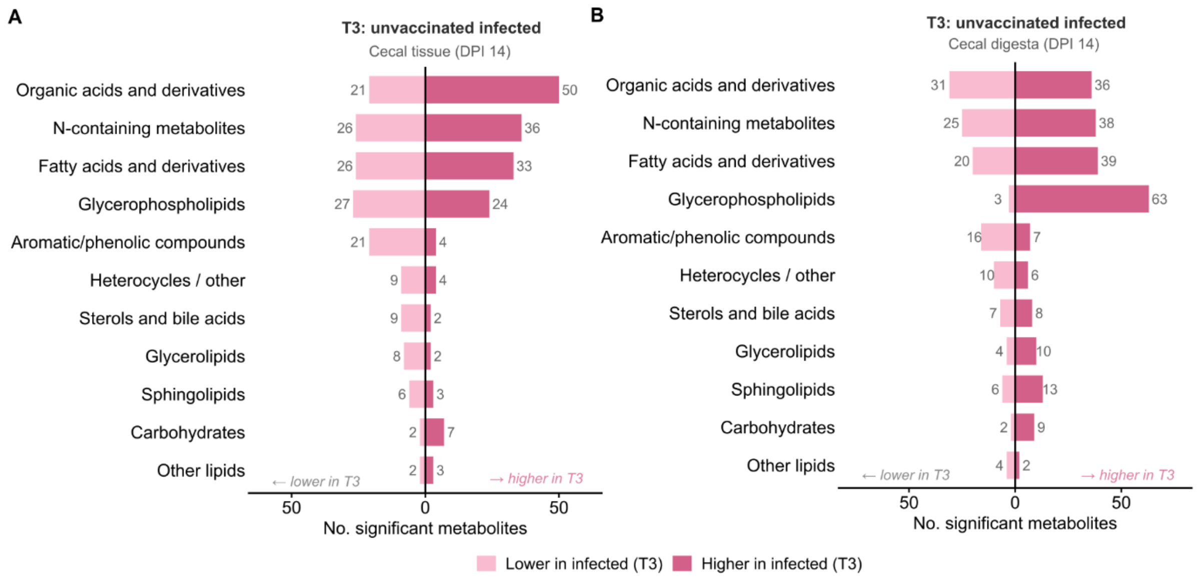


**Fig. S3. Number of significantly altered identified metabolites per chemical superclass in unvaccinated infected birds (T3) versus the negative control (T4) at DPI 14 in cecal tissue (A) and digesta (B)**. Dark bars indicate metabolites elevated in T3; light bars indicate metabolites reduced in T3. Analysis is based on n = 555 uniquely identified metabolites spanning 11 chemical superclasses (adj.p $< 0.05$).

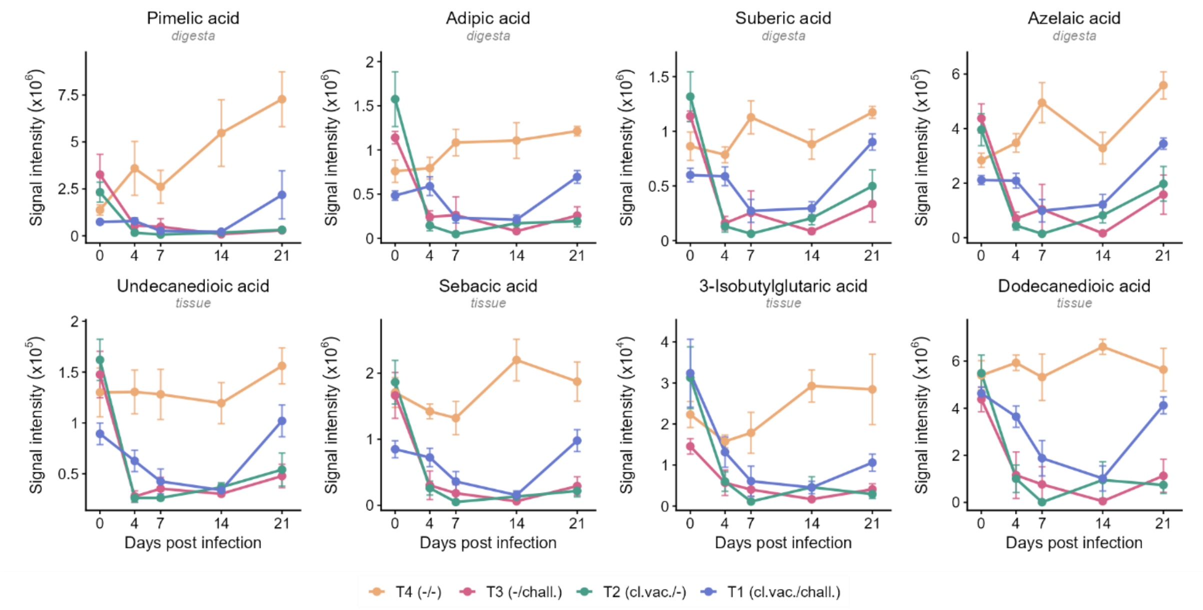


**Fig. S4. Extended dicarboxylic acid and branched-chain catabolite profiles**. Temporal profiles of pimelic acid (digesta), adipic acid (digesta), suberic acid (digesta), azelaic acid (digesta), undecanedioic acid (tissue), sebacic acid (tissue), 3-isobutylglutaric acid (tissue), and dodecanedioic acid (tissue). Data are presented as mean ± SEM. T1 (cl.vac./chall.): cloacal vaccination followed by challenge; T2 (or.vac./chall.): oral vaccination followed by challenge; T3 (-/chall.): challenged only; T4 (-/-): unvaccinated unchallenged control.

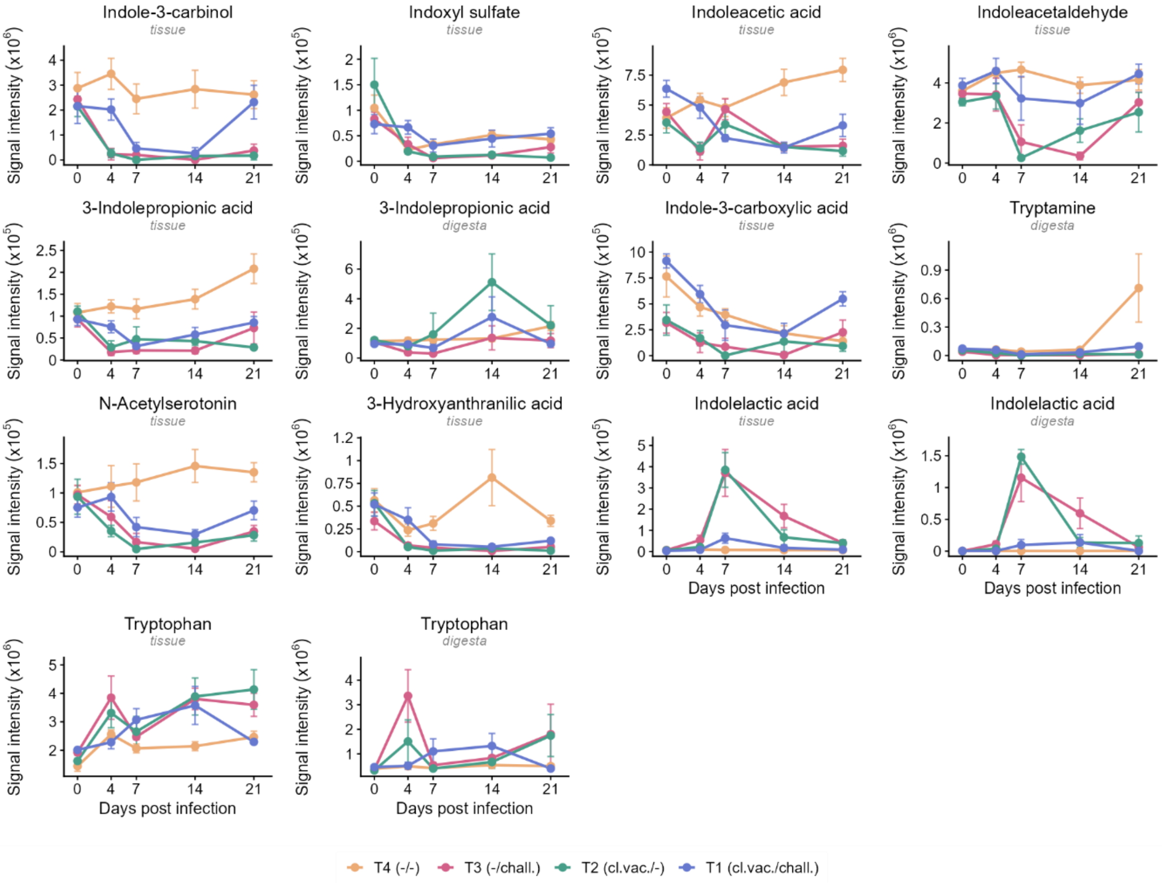


**Fig. S5. Extended tryptophan and indole pathway metabolite profiles.** Temporal profiles of indole-3-carbinol (tissue), indoxyl sulfate (tissue), indoleacetic acid (tissue), indoleacetaldehyde (tissue), 3-indolepropionic acid (tissue and digesta), indole-3-carboxylic acid (tissue), tryptamine (digesta), N-acetylserotonin (tissue), 3-hydroxyanthranilic acid (tissue), indolelactic acid (tissue and digesta), and tryptophan (tissue and digesta). Note that indolelactic acid was elevated during active infection, in contrast to all other indole metabolites shown. Data are presented as mean ± SEM. T1 (cl.vac./chall.): cloacal vaccination followed by challenge; T2 (or.vac./chall.): oral vaccination followed by challenge; T3 (-/chall.): challenged only; T4 (-/-): unvaccinated unchallenged control.

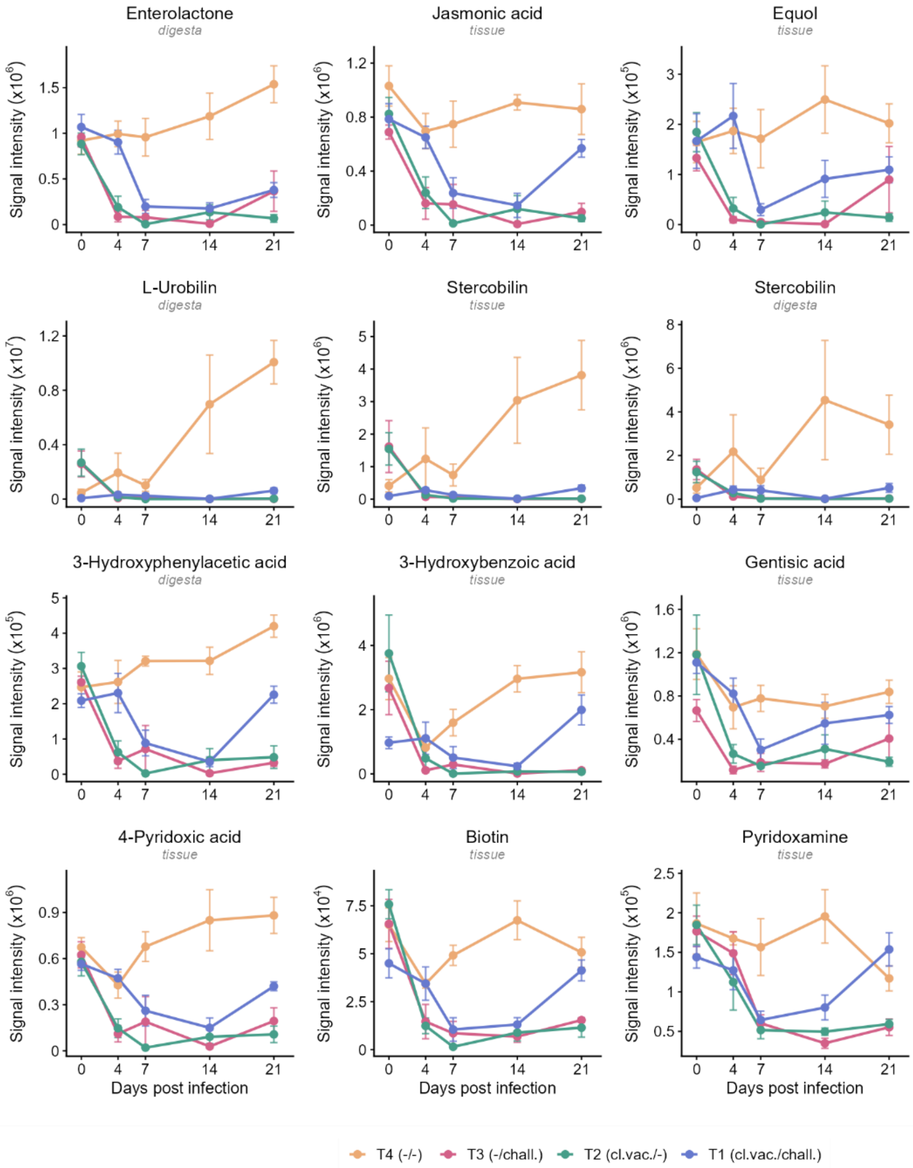


**Fig. S6. Extended phytochemical biotransformation, phenolic catabolite, bilirubin catabolites, and vitamin-related metabolite profiles**. Temporal profiles of enterolactone (digesta), jasmonic acid (tissue), equol (tissue), urobilin (digesta), stercobilin (tissue, digesta), 3-hydroxyphenylacetic acid (digesta), 3-hydroxybenzoic acid (tissue), gentisic acid (tissue), 4-pyridoxic acid (tissue), biotin (tissue), and pyridoxamine (tissue). Data are presented as mean ± SEM. T1 (cl.vac./chall.): cloacal vaccination followed by challenge; T2 (or.vac./chall.): oral vaccination followed by challenge; T3 (-/chall.): challenged only; T4 (-/-): unvaccinated unchallenged control.

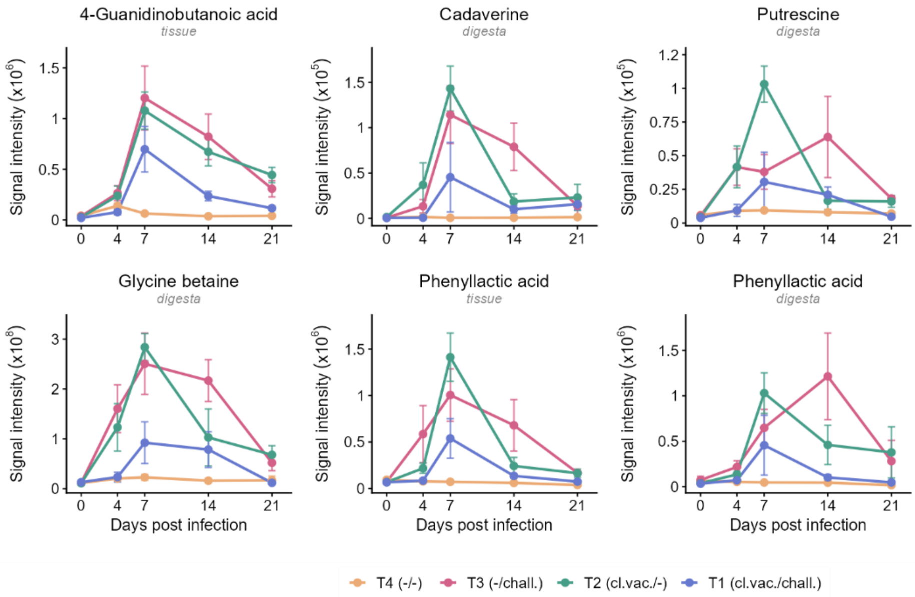


**Fig. S7**. **Extended accumulating metabolite profiles**. Temporal profiles of 4-guanidinobutanoic acid (tissue), cadaverine (digesta), putrescine (digesta), glycine betaine (digesta), phenyllactic acid (tissue and digesta). Data are presented as mean ± SEM. T1 (cl.vac./chall.): cloacal vaccination followed by challenge; T2 (or.vac./chall.): oral vaccination followed by challenge; T3 (-/chall.): challenged only; T4 (-/-): unvaccinated unchallenged control.

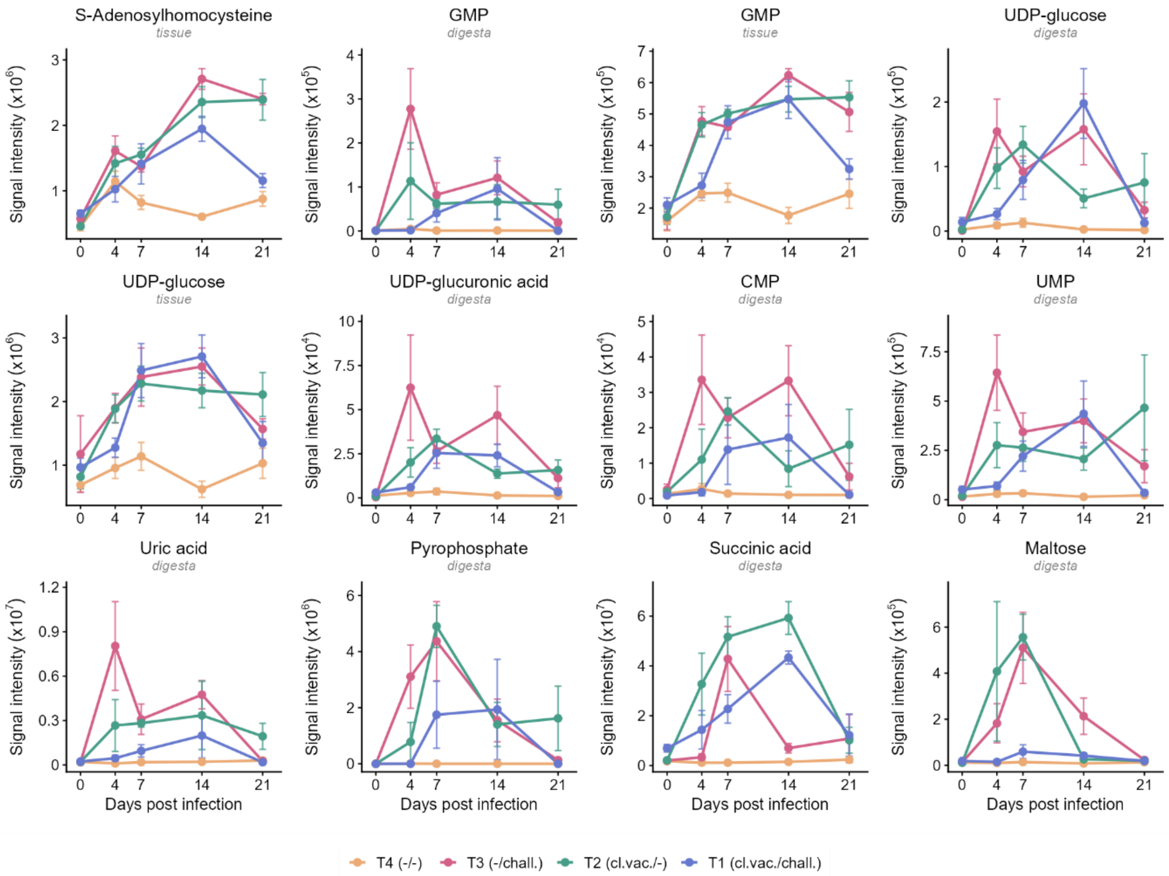


**Fig. S8**. **Extended inflammatory activation and nucleotide accumulation profiles**. Temporal profiles of S-adenosylhomocysteine (SAH, tissue), GMP (digesta and tissue), UDP-glucose (digesta and tissue), UDP-glucuronic acid (digesta), CMP (digesta), UMP (digesta), uric acid (digesta), pyrophosphate (digesta), succinate (digesta), and maltose (digesta). Data are presented as mean ± SEM. T1 (cl.vac./chall.): cloacal vaccination followed by challenge; T2 (or.vac./chall.): oral vaccination followed by challenge; T3 (-/chall.): challenged only; T4 (-/-): unvaccinated unchallenged control.

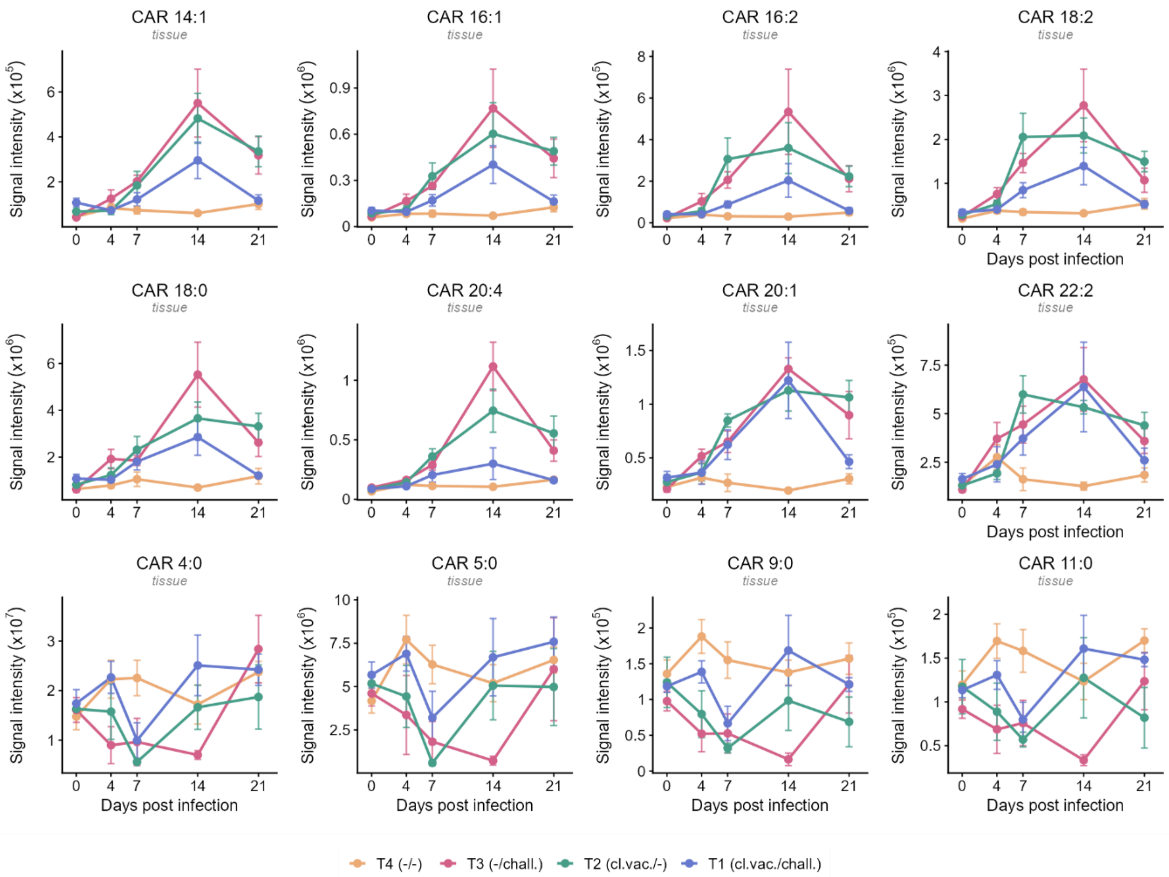


**Fig. S9**. **Extended acylcarnitine profiles**. Temporal profiles of elevated long-chain acylcarnitines CAR 14:1, CAR 16:1, CAR 16:2, CAR 18:2, CAR 18:0, CAR 20:4, CAR 20:1, and CAR 22:2 (tissue), and depleted short-to-medium chain acylcarnitines CAR 5:0, CAR 9:0, and CAR 11:0 (tissue). Data are presented as mean ± SEM. T1 (cl.vac./chall.): cloacal vaccination followed by challenge; T2 (or.vac./chall.): oral vaccination followed by challenge; T3 (-/chall.): challenged only; T4 (-/-): unvaccinated unchallenged control.

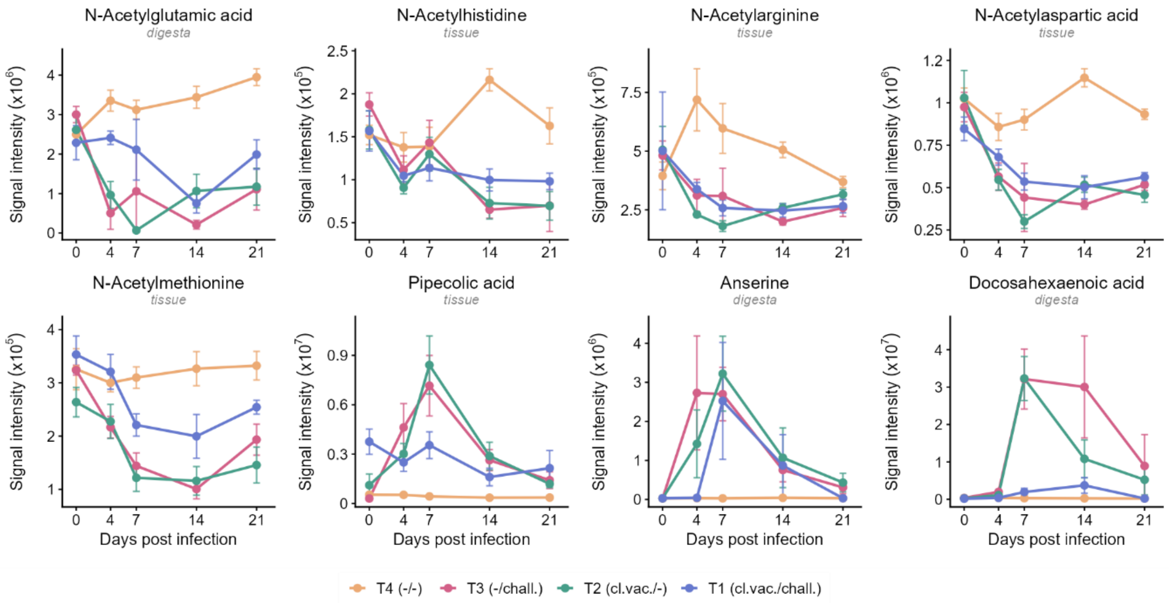


**Fig. S10**. **Extended N-acetylated amino acid, pipecolic acid, anserine, and docosahexaenoic acid profiles**. Temporal profiles of N-acetylglutamic acid (digesta), N-acetylhistidine, N-acetylarginine, N-acetylaspartic acid, and N-acetylmethionine (tissue), pipecolic acid (tissue), anserine (digesta), and docosahexaenoic acid (digesta). Data are presented as mean ± SEM. T1 (cl.vac./chall.): cloacal vaccination followed by challenge; T2 (or.vac./chall.): oral vaccination followed by challenge; T3 (-/chall.): challenged only; T4 (-/-): unvaccinated unchallenged control.

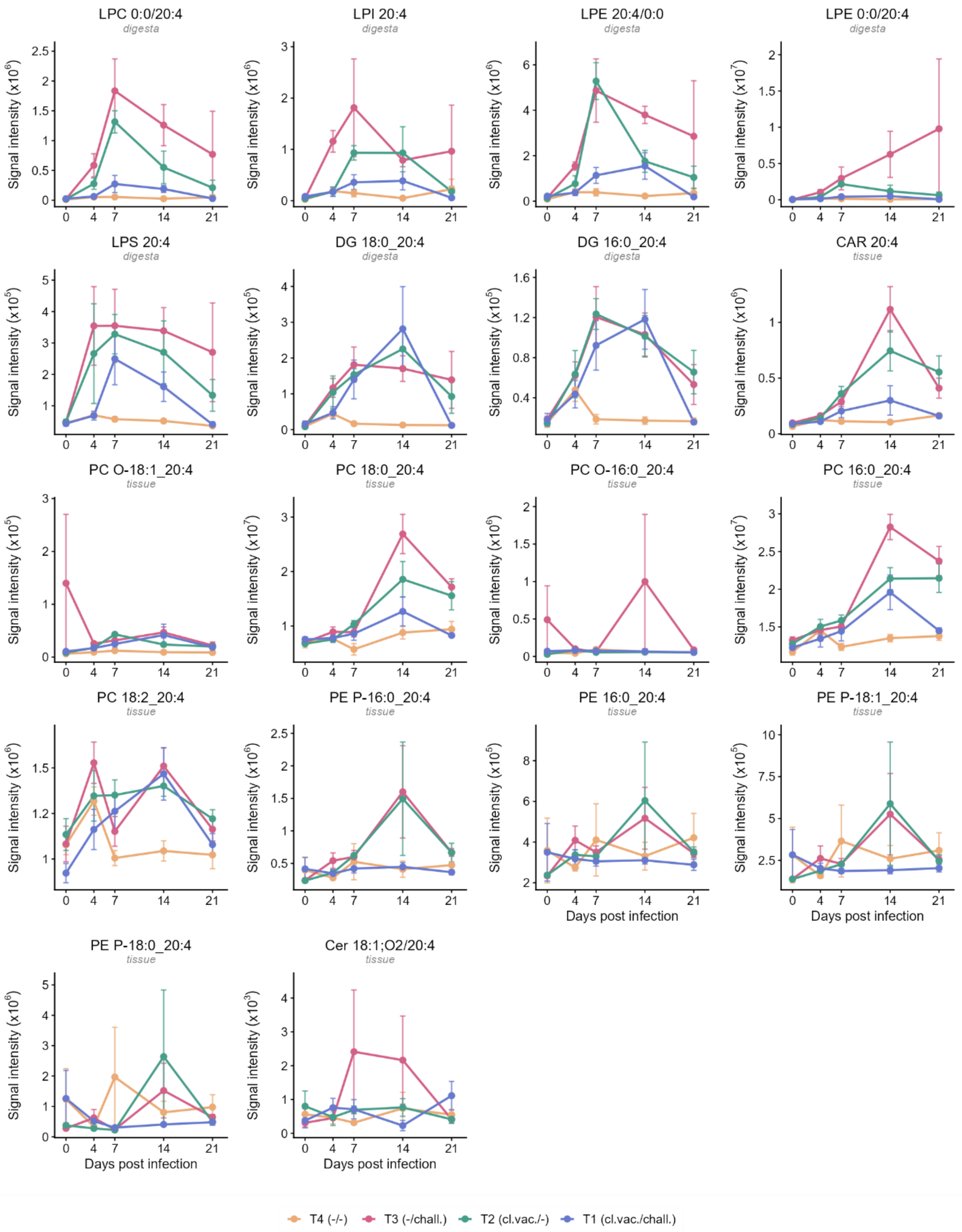


**Fig. S11**. **Extended Arachidonic acid-containing lipid profiles during infection.** Temporal profiles of lysophospholipids (LPC 0:0/20:4, LPI 20:4, LPE 20:4/0:0, LPE 0:0/20:4, LPS 20:4; digesta), diacylglycerols (DG 18:0_20:4, DG 16:0_20:4; digesta), arachidonoylcarnitine (CAR 20:4; tissue), phosphatidylcholines (PC O-18:1_20:4, PC 18:0_20:4, PC O-16:0_20:4, PC 16:0_20:4, PC 18:2_20:4; tissue), phosphatidylethanolamines (PE P-16:0_20:4, PE 16:0_20:4, PE P-18:1_20:4, PE P-18:0_20:4; tissue), and ceramide (Cer 18:1;O2/20:4; tissue). Data are presented as mean ± SEM. T1 (cl.vac./chall.): cloacal vaccination followed by challenge; T2 (or.vac./chall.): oral vaccination followed by challenge; T3 (-/chall.): challenged only; T4 (-/-): unvaccinated unchallenged control. LPC, lysophosphatidylcholine; LPI, lysophosphatidylinositol; LPE, lysophosphatidylethanolamine; LPS, lysophosphatidylserine; DG, diacylglycerol; CAR, acylcarnitine; PC, phosphatidylcholine; PE, phosphatidylethanolamine; Cer, ceramide; P-, plasmanyl; O-, plasmenyl.

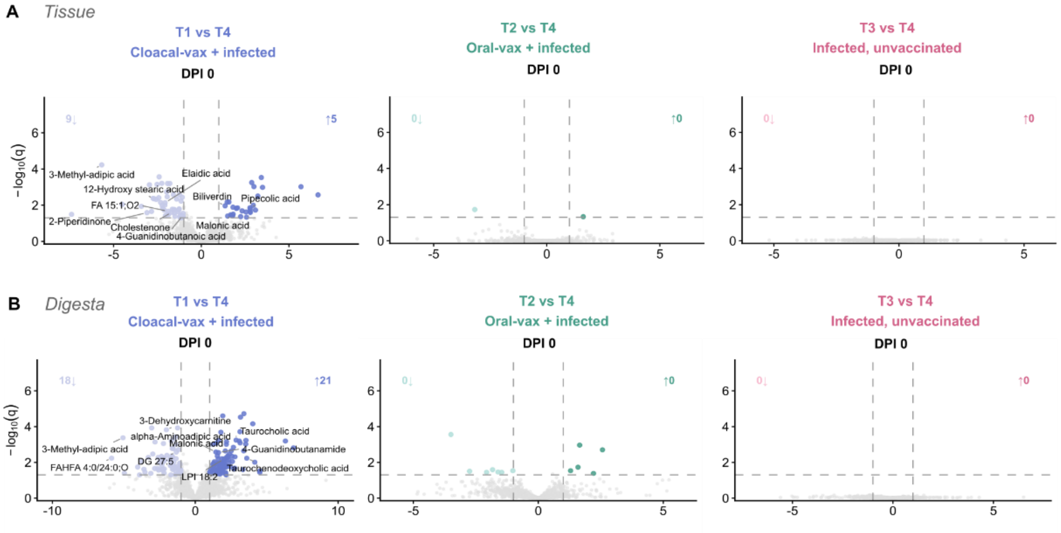


**Fig. S12. Cloacally vaccinated birds showed a distinct pre-challenge metabolic shift**. Volcano plots showing significantly altered identified metabolites in cecal tissue for each treatment group versus the negative control (T4) before *H. meleagridis* challenge (DPI 0) in cecal tissue **(A)** and digesta **(B)**. Columns correspond to treatment groups: T1 (cloacal vaccination/challenge, blue), T2 (oral vaccination/challenge, green), and T3 (challenge only, unvaccinated, pink) when compared to negative control (T4). Each point represents one identified metabolite; colored points passed both significance (adj.p (q) < 0.05, Benjamini-Hochberg correction) and effect size ($|log_2FC| \geq 1$) thresholds. Corner numbers indicate the counts of identified metabolites passing both thresholds. The top 10 metabolites were labeled in the volcano plots.
